# Stabilization of extensive fine-scale diversity by spatio-temporal chaos

**DOI:** 10.1101/736215

**Authors:** Michael Pearce, Atish Agarwala, Daniel S. Fisher

## Abstract

It has become recently apparent that the diversity of microbial life extends far below species level to the finest scales of genetic differences. Remarkably, extensive fine-scale diversity can coexist spatially. How is this diversity stable on long timescales despite selective or ecological differences and other evolutionary processes? Most work has focused on stable coexistence or assumed ecological neutrality. We present an alternative: extensive diversity maintained by ecologically-driven spatio-temporal chaos, with no assumptions about niches or other specialist differences between strains. We study generalized Lotka-Volterra models with anti-symmetric correlations in the interactions inspired by multiple pathogen strains infecting multiple host strains. Generally, these exhibit chaos with increasingly wild population fluctuations driving extinctions. But the simplest spatial structure, many identical islands with migration between them, stabilizes a diverse chaotic state. Some types (sub-species) go globally extinct, but many persist for times exponentially long in the number of islands. All persistent types have episodic local blooms to high abundance, crucial for their persistence as, for many, their average population growth rate is negative. Snapshots of the distribution of abundances show a power-law at intermediate abundances that is essentially indistinguishable from the neutral theory. But the dynamics of the large populations are much faster than birth-death fluctuations. We argue that this spatio-temporally chaotic “phase” should exist in a wide range of models, and that even in rapidly mixed systems, longer lived spores could similarly stabilize a diverse chaotic phase.

## 1 Introduction

Enormous diversity of species is one of the remarkable features of life on Earth. Once it has been established by evolution, this diversity is traditionally explained in terms of niches and geographical separation. But even then, spatial coexistence of a wide variety of species that seem to occupy similar niches is a major puzzle [1].

Recent studies have found that even within individual microbial species (e.g. *Vibrio* [2] in the ocean, *Synechococcus* in hot springs [3, 4], *S. epidermis* on the skin [5, 6], *Neisseria* on the tongue [7] and *B. vultatus* in human guts [8]), many strains differing genetically on a broad spectrum of scales can coexist in the same spatial location or nearby. This fine-scale (or micro-) diversity is especially surprising when strains mix together and are hence forced to compete. In some cases, the relevant mixing times are known: for the most abundant phytoplankton species, *Prochlorococcus*, which dominates tropical mid-oceans [9], a single sample contains hundreds of different strains which diverged over timescales much longer than mixing times of the ocean [10]. Very similar sub-types are found in both the Atlantic and Pacific [11].

Why doesn’t survival of the fittest eliminate fine-scale diversity on time scales that are long compared to generation or spatial mixing times, but still short compared to the evolutionary time scales over which the diversity must have evolved and be maintained? To understand this, is it necessary to interpret the strains, sub-strains, and sub-sub-strains as “ecotypes” [12] adapted to microniches and differing phenotypically in essential ways? Or might there be more general explanations? Any satisfying theory should lead to understanding of how the statistical structure of diversity — not just its existence — arises and is maintained by evolution.

Microbial diversity is often characterized by abundance distributions. Abundances typically range over many decades extending down to very rare species, and are often fit by power-laws [13, 14, 15]. However data on abundance distribution of within-species fine-scale diversity is still rather limited [3, 4, 7]. Furthermore, dynamical data is crucial to distinguish scenarios — as we shall discuss – but is much harder to come by due to the need for deeply sequenced time-series data, as in, e.g. [14, 15, 16]. Is local fine-scale diversity relatively stable? Or are blooms from low to high abundance and back down — observed in a variety of contexts [17, 18] — more the norm?

Theoretical ecologists have long endeavored to discover the ingredients necessary for diversity to persist in complex ecosystems. Most of this work has focused on *ecologically stable diversity* (reviewed in [19]). Common approaches include modeling interactions via competition for resources [20, 21, 22, 23, 24], or approximating inter-species interactions as “random” [25, 26, 27, 28, 29, 30]. A standard assumption is that species, or strains, compete with themselves, and hence suppress their own growth, more strongly than they interact with other types: this is equivalent to assuming that each species has its own niche [27, 28, 29, 31, 32]. Niches may be a good starring point for modeling interactions among different species. But for closely related strains there is no obvious reason why the interactions between “siblings” should be much stronger than between distant “cousins”.

The opposite extreme to niche-based approaches is the *neutral theory of biodiversity* [33, 34] which posits that broad classes of species – such as all trees in some region [35] or all diatoms of similar size [36] – are ecologically equivalent. Species abundances fluctuate due to neutral birth, death, and migration processes instead of being stabilized by ecological interactions. The balance between these results in a broad distribution of abundances with a simple power law tail [37, 38] that compares well, at least semi-quantitatively, to data on a variety of systems [39, 40, 41].

But for microbial diversity, there is a major quantitative problem with neutral theory: with the large sizes of microbial populations, birth-death fluctuations are far too slow to dominate population dynamics. The short generation times mean that even tiny selective differences will be greatly amplified on modest time scales [42]. To produce broad distributions from faster dynamics, recent work has generalized effective neutrality by considering species with different responses to a fluctuating environment but still neutral time-averaged fitnesses [43, 44]. But this does not solve the problem of fluctuating to extinction, nor of how the average neutrality might arise.

A different approach, which we take here, is to ask whether ecological interactions and fitness differences can drive continually changing abundances in a state of *chaotic coexistence*. As recognized early by May [45, 46], a range of simple deterministic ecological models exhibit abundances that vary chaotically [47]. It has been proposed that dynamical coexistence is important for plankton biodiversity [48, 49] and chaos has been demonstrated in a long term experiment of an isolated planktonic food web [50]. In nature, rapid changes in viral and microbial strains have been observed in controlled, aquatic environments [51] and cycling between abundant and rare is commonly observed in oceanic bacterial taxa [52]. Might chaos be a generic feature of complex ecosystems [53, 54]? Can chaos promote or stabilize extensive diversity among close relatives?

A ubiquitous driving force of evolution is the advantage of evolving to *“kill-the-winner”* — predation, pathogenicity, or arms races with relatives such as among *Streptoomyces* bacteria [55]. For pathogens, increasing abundance of a host strain can result in increasing abundance of particular pathogen strains that limit the successful host population and subsequently those pathogen strains as well.

The original predator-prey Lotka-Volterra (LV) model demonstrated dynamical coexistence with periodic variations of the predator and prey populations [56, 57, 58]. What about kill-the-winner dynamics with many interacting populations? Models with specialist pathogens can lead to stable static communities [59, 60], as indeed does slight modification of the original LV model. Coupled predator-prey cycles have been studied [61, 62]; generically, multiple coupled cycles tend to lead to chaos.

A major drawback of most models of host-pathogen dynamics, is that they do not account for broadly non-specialist pathogens — analogous to assuming niches. This may be reasonable for multi-species communities, but it is not for understanding within-species fine-scale diversity. Indeed, phages are often found to infect related but phenotypically distinct bacterial strains [63, 64]. A crucial need for understanding fine scale diversity is models with many strains of a single host species and of a single pathogen species, with broad infectivity – but its strength varying between strains.

For microbial populations, evolutionary and ecological time scales overlap and diversity is always being produced. However there is an important question of principle — as well as, quantitatively, in practice: Does evolution generally lead to extensive spatially-coexisting fine-scale diversity that persists for much longer than ecological time scales? Or does evolution tend to destroy diversity of communities assembled from previously isolated strains? A number of studies of simple models of stable coexistence, show that the diversity tends to collapse when processes like mutation or invasion of new types are included [65]. However this depends on how the evolution is modeled and others have reached the opposite conclusion [66, 67].

What are the consequences of evolution in host-pathogen models? Although it is often said that pathogens promote diversity [68], this has not been convincingly demonstrated in general models. Recent work has shown that if evolution is fast enough, extensive chaotic diversity of host-pathogen systems can evolve and persist [69]. Others have explored this by putting in artificial forms of the population dynamics that save multiple types from extinction [70], but one needs to go beyond this. Thus we focus here on a key underlying question: Can generalized kill-the-winner-like ecological interactions, without any special assumptions, stabilize extensive diversity on long time scales?

We analyze a general class of Lotka-Volterra models and investigate the nature of a diverse chaotic state that exists in a special case [71]. More generically, we show that the chaos drives a cascade of extinctions and destroys the diversity. But the simplest form of spatial structure — identical islands with migration between them — leads to a stable spatio-temporally chaotic “phase” that we argue should occur far more generally. We conclude with open questions for future work, especially about the effects of evolution.

## 2 Models of complex diverse ecosystems

We focus on generalized Lotka-Volterra (LV) models widely used to model ecological interactions between species. For competing species, the dynamics of the population size, *n*_*i*_, of species *i* is conventionally written in terms of a basic population growth rate, *r*_*i*_, a carrying capacity *κ*_*i*_ and a matrix *W*_*ij*_ of interactions between them:

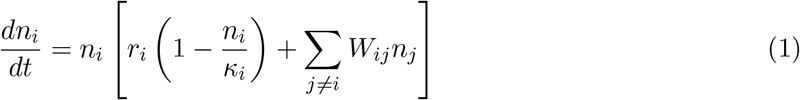

This implicitly assumes that each species has its own “niche” and treats the interactions between species differently than interactions within species.

For closely related subtypes, our focus, there is no *a priori* reason that interactions with the same type is particularly different than with other types. It is useful to separate out the overall constraints from use of the same resources which will keep the total population, 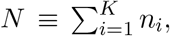, approximately constant, and focus on the x differences. We then write dynamics in terms of the *frequencies*, *ν*_*i*_ ≡ *n*_*i*_/*N*

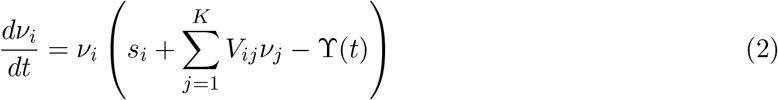

where 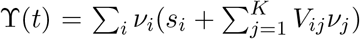 keeps the total number fixed (Σ_*i*_*ν*_*i*_ = 1). The interactions, including with the same type, can be considered with reference to a (fictitious) “average” type labeled 0, as *V*_*ij*_ = *N*[*W*_*ij*_ − *W*_*i*0_ − *W*_0*j*_ + *W*_00_]. The overall selective differences, *s*_*i*_ = *r*_*i*_ − *r*_0_ + [*W*_*i*0_ − *W*_00_]*N*, include a part representing how type *i* does against the “average” of the others.

Competition for resources will result in positive correlations between how types interact with each other, e.g.. between *V*_12_ and *V*_21_, while direct one-on-one competition will result in anticorrelations: if type 1 “beats” type 2 so that *V*_12_ > 0, then 2 “loses” to 1 so that *V*_21_ < 0. Antisymmetric correlations of the interactions also occur for systems of hosts and pathogens in which a collection of strains of a pathogen infect a spectrum of strains of a host – e.g. intra-species diversity within one phage and one bacterial species [72, 73]. In terms of a population vector, 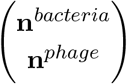, the interaction matrix has a block diagonal structure 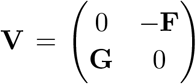. For closely related types, **F** and **G**^*T*^ will be positively correlated since interactions that are better for the phage are worse for the bacteria and visa versa.

For simplicity we will focus on interactions between strains of a single species. Being sums and differences between similar interactions and similar growth rates, it is then natural to treat the *V*_*ij*_ and *s*_*i*_ as approximately random and characterize them by their statistical properties. The magnitudes of the *V*_*ij*_ set the time scale of the ecological dynamics. For most of our analyses, we will assume that the effects of overall fitness differences, the {*s*_*I*_}, are much smaller than the differences in the influences of the interactions. (This corresponds to the assumption that generalists are hard to evolve.) The simplest choice for the distribution of the interactions is that the {*V*_*ij*_}, for *i* ≠ *j*, are Gaussian distributed with mean zero, identical variance, and all covariances zero except across the diagonal. The ratio

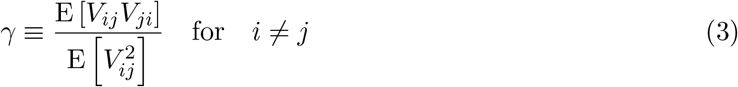

with −1 ≤ γ ≤ 1, characterizes the correlation between how two types affect each other: γ = −1 corresponds to an antisymmetric matrix, γ = 0 to independent elements, and γ = 1 to a symmetric >matrix. If, in spite of the similarities between types, competition with others of the same type are stronger — niche-like interactions — the diagonal terms would be negative on average and we thus define 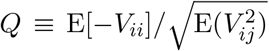. An important point, however, is that for a large number of interacting types the niche-like interactions must be much stronger than the interactions between different types: as the sum over the (random) effects of all the other types will be of order 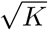, in order to substantially affect the ecological dynamics, *Q* must be of order 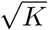. Most theoretical work, motivated particularly by competition for a mixture of resources [20, 21, 48, 74], has focused on such large *Q* [27, 28, 75, 76] and positive correlations between interactions, γ > 0 [23, 24, 29, 30, 31, 32]. In this regime, for a large number of types, there is a unique stable, uninvadable, community, corresponding to a stable fixed point of the dynamics, with a substantial fraction 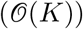 types surviving and the other types unable to invade. This occurs, for any γ, when 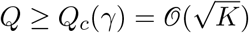. For smaller *Q* the fixed point destabilizes and it appears that there is generally no large stable community (except for the special case of γ = 1 [71, 76]). As we are interested in the interaction between large numbers of similar types, we will mostly neglect the effects of niche-like interactions and set *Q* = 0.

What happens when there is no large stable community? The original Lotka-Volterra of an interacting predator and prey is a special case with *Q* = 0 and γ = −1: it exhibits a family of periodic orbits [56, 57]. Our focus will primarily be on γ < 0 for which dynamical ecologies are ubiquitous as has been found in [58, 71, 76]. We first analyze the special case γ = −1. As we shall see, many aspects of the chaotic dynamics in this limit well characterize the behavior for a much broader range of parameters and models and the analysis of this special case serves as a needed framework for understanding chaotic dynamics more generally including, crucially, the effects of spatial structure.

## 3 Chaotic Ecological Dynamics

### 3.1 Perfectly antisymmetric Lotka-Volterra

The idealized model of perfectly *antisymmetric* interactions: *V*_*ij*_ = −*V*_*ji*_., has, in the deterministic limit of infinite population size, special properties. We first summarize its key properties — illustrated in Figures 1 and 3 — and then outline their derivation.

**Figure 1:**
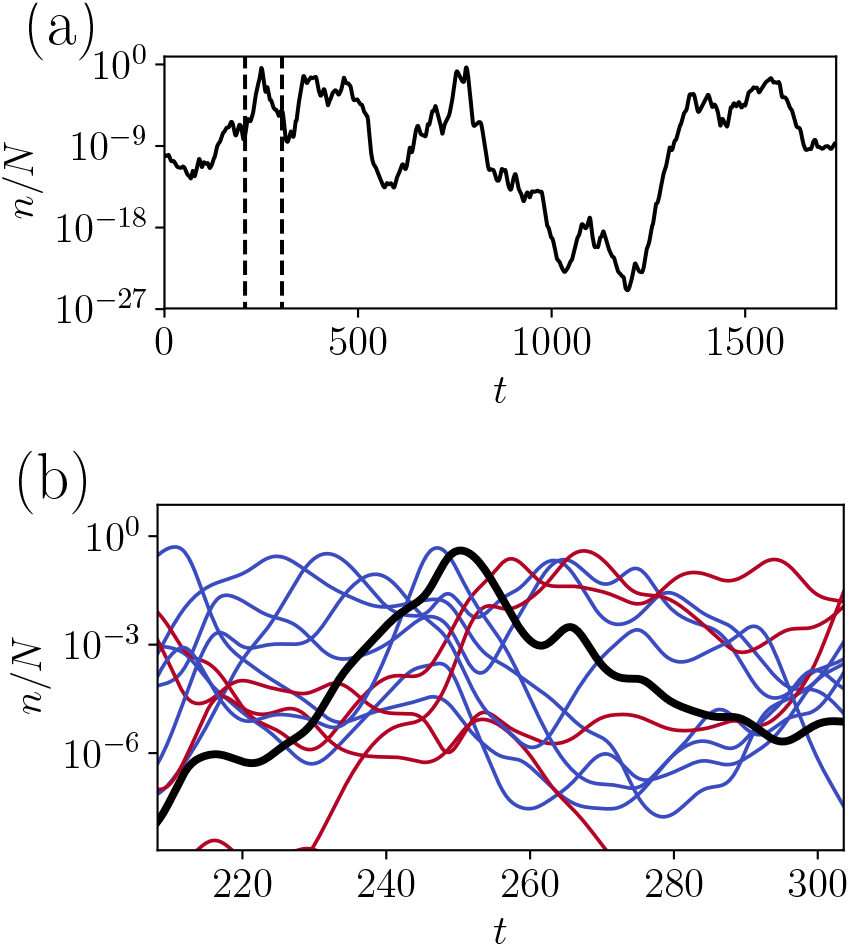
*Chaotic dynamics of the perfectly antisymmetric model (ASM) Top*: Population dynamics of the frequency (fractional abundance), *ν*_*i*_(*t*) = *n*_*i*_(*t*)*/N*, of a single type, out of a total of *K* = 301 persistent types (of 600 initial types). The range on the logarithmic scale is set by the “temperature”, Θ, here 40 (unrealistically large to emphasize the different time scales). Log(*ν*_*i*_) exhibits super-diffusive random-walk like behavior over a wide-range of time scales. Note the multiple peaks that occur before *ν* fluctuates down to low abundance which typically takes time 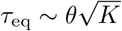. *Bottom*: Short time dynamics near a high-abundance peak of type *i* (black). Its rise is due to the high abundance of several other types, *j*, (blue) with positive effects on type *i* via *V*_*ij*_*ν*_*j*_ > 0. The high abundance of *i* then brings up populations of some other types, *k*, red. Because of the antisymmetry, the effects of red types on the black type, via *V*_*ik*_*ν*_*k*_ < 0, cause negative feedback that drives the black population back down. Individual peaks typically last for short times 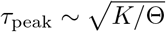.

The antisymmetric model (ASM) has a family of stable chaotic states in which a fraction of the types are extinct and cannot invade while the rest — a unique set — coexist in a chaotic steady-state. Because the effects of the interactions make the growth or decay of the population vary, the logarithms of the frequencies, log(*ν*_*i*_(*t*)) are the natural variables. The log(*ν*_*i*_) of the persistent types fluctuate over some range of width Θ; this parameter, which turns out to be analogous to a temperature, parameterizes the family of chaotic steady states. We concentrate on large Θ for reasons that will become apparent later. While the frequency of each type fluctuates wildly, each will sometimes reach high frequency and when it does so will have a large effect on types that do better in its presence. But as these others rise to high frequency themselves, the negative effect that they have on the first type — implied by the antisymmetry — will cause its frequency to peak and start decaying as illustrated in Figure 1(b). These peaks typically last for a short time, 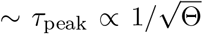. Although they only occur rarely, they will dominate the average frequency of each type, ⟨*ν*_*i*_⟩, which will vary from type-to-type.

Over a much longer time scale, *τ*_eq_ ∝ Θ, log(*ν*_*i*_(*t*)) fluctuates over its full range, but on intermediate time scales, it undergoes a super-diffusive random walk, seen in Figure 1(a), whose statistical properties will turn out to be crucial and much more general than the ASM. We now derive these results, with some details relegated to Appendix B.

#### Static properties of the antisymmetric model

The ASM has several special properties. The average over all interactions Σ_*i,j*_ *V*_*ij*_*ν*_*i*_*ν*_*j*_ is always 0. Thus the total number is conserved without the Lagrange multiplier term: ϒ = 0. It can be shown that for antisymmetric *V*_*ij*_, there exists a *unique uninvadable fixed point* 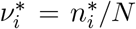. At this fixed point some types coexist, with 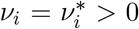, set by 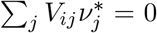, for *i* in the coexisting subset, while others go *extinct*, with 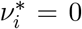 and negative average growth rate, 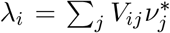 so they cannot invade if seeded at small numbers (indeed, for this special model, not at all) [76].

Away from the fixed point, there is a Lyapunov function of the dynamics which is determined by the uninvadable fixed point:

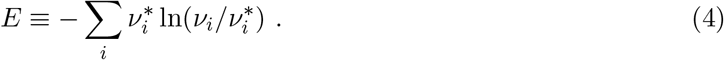

This is strictly decreasing as long as there are types destined for extinction. But when these have gone extinct, *E* is conserved. There thus exists a one-parameter family of “states” with *E* the weighted distance, in log-space, between the current set of frequencies and the fixed point set of frequencies. The conservation of *E* ensures that all the *persistent* (non-extinct) types coexist indefinitely.

When only these persistent types survive, the phase space density (volume form) of their log frequencies, {*x*_*i*_} ≡ {log(*ν*_*i*_)} of these types, is conserved. If the dynamics is ergodic, as expected in the limit of large *K*, the probability distribution of the frequencies of the persistent subset is the usual microcanonical ensemble with energy *E* and total number. For large *K*, this is essentially equivalent to the grand canonical ensemble of *independent* — but not equivalent — frequencies as noted in [71]:

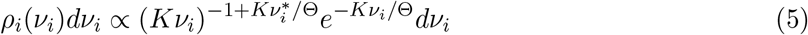

with the chemical potential-like term (= −Θ) chosen to give, on average, Σ_*i*_ *ν*_*i*_ = 1. A simple computations gives ⟨E⟩ = Θ. The form of the distribution suggests that only of order *K*/Θ types dominate at any given time. For this distribution 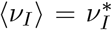 for each *i*, independent of Θ, where ⟨·⟩ is a time-average (equivalent to the average over this ensemble). For large Θ, the population of each type is distributed roughly uniformly on a log scale up to a soft “ceiling” ~ Θ/*K*, independent of *i*, and exponentially suppressed at large negative 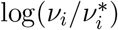 of order Θ.

Intriguingly, the form of Equation 5, as well as the joint distribution of all the {*ν*_*i*_} with fixed total population size (for which the exponential factors are replaced by *δ*(Σ_*i*_ *ν*_*i*_ − 1)) is *identical* to the frequency distribution from *neutral* ecological dynamics with genetic drift and random migration from a “mainland” (with immigrants arriving at a rate 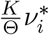 for type *i* [77, 78]). However, as we describe in the next section, the deterministic chaotic dynamics is quite different than stochastic neutral dynamics.

### 3.2 Dynamical mean-field theory

Even for the special ASM, the dynamics cannot be found exactly. However, the dynamics for this, as well as the much more general models we will study, can be analyzed by dynamical mean-field theory (DMFT) [79]. DMFT directly applies when *V*_*ij*_ is a large random matrix — in our case with off-diagonal elements i.i.d. except for the covariance parameterized by γ (Equation 3). This approach approximates the deterministic driving of each type by many others with a stochastic integro-differential equation with a type driven by a combination of a Gaussian noise from all the other types, in its absence, together with a feedback from the effect of that type on all the others and the resulting effect back on it. The dynamical mean-field is asymptotically exact when, at all times, the dynamics of each types is influenced by a large number of others: I.e., *K* → ∞.

With the variances of the *V*’s (chosen to be unity) setting the time scale, the dynamics of type *i* is

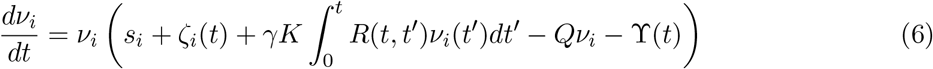

We will refer to this definition of the dynamics as the “natural” scaling. Here *ζ*_*i*_(*t*) represents the effects of all the others on the growth-minus-death rate of type *i* when *ν*_*i*_ is small: *ζ*_*i*_(*t*) is Gaussian with covariance self-consistently determined by

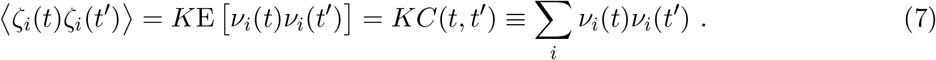

where the E denotes averages taken over the full ensemble (that is, the distribution the *V*_*ij*_ induce on trajectories). The *ζ*_*i*_ for different *i* are independent for *K* → ∞ (the simplification that makes dynamical mean-field theory exact). The second term in Equation 6 represents the effects of *V*_*ji*_*ν*_*i*_(*t*′) on 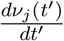 and the resulting feedback at a later time on *ν*_*i*_(*t*) via Σ_*j*_ *V*_*ij*_*ν*_*j*_(*t*), so that *R* must be the

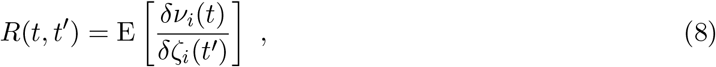

whose strength of action is parameterized by γ.

Unfortunately, in contrast to many studied problems, there are no closed equations for the response and correlation functions so we must analyze the stochastic problem directly. Studying fixed points is much simpler [28, 29, 80]; a primary technical contribution of this work is the first quantitative analysis of the dynamics and its broader consequences.

#### Antisymmetric model mean-field theory

We first analyze the ASM, γ = −1, with *Q* = 0, and all *s*_*i*_ = 0 building on the exact results for the probability distributions of the {*ν*_*i*_} in the ASM. (In Appendix D.3 we will analyze the quantitative modifications due to selective differences, which can be included by adding *s*_*i*_ − *s*_*j*_ to *V*_*ij*_, keeping it antisymmetric.) Over the ensemble of random antisymmetric matrices with i.i.d. elements, it can be shown [76] that every subset with an odd number of types has equal chance of surviving at the unique uninvadable fixed point so that for large *K*, 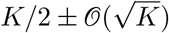 types survive. The uninvadable fixed point corresponds to Θ = 0.

For any Θ > 0, the behavior is chaotic with the persistent types undergoing fluctuations in log-frequencies *x*_*i*_ ≡ log(*ν*_*i*_). (Figure 1). By Equation 5 the instantaneous (−*x*_*i*_) are exponentially distributed over a scale 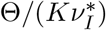. The key to understanding the dynamics for large Θ lies in the statistics of the peaks (Figure 1) - the times when a type fluctuates to its largest values which are *ν*_*i*_ ∼ Θ/*K*,*independent* of 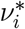. Thus Θ also parameterizes the heights of the peaks: each population will fluctuate up to Θ times the average population per type. At any time, a fraction of order 1/Θ of the types will be peaking and these will dominate the forces on the others: thus for DMFT to be valid requires *K*/Θ ≫ 1 and we restrict consideration to this regime.

In a statistical steady state, *C*(*t, t*′) = *C*(*t*−*t*′) is a function of time differences only and similarly *R*(*t, t*′) = *R*(*t* − *t*′). At long times *C*(*t* − *t*′ → ∞) attains the non-zero value 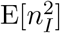. Thus the noise *ζ*_*i*_ has a type-dependent average “bias” *ξ*_*i*_, with zero mean and variance *KC*(*t* − *t*′ → ∞). One can thus rewrite *ζ*_*i*_(*t*) = *ξ*_*i*_ + *η*_*i*_(*t*), where *η*_*i*_(*t*) is the fluctuating part of *ζ*_*i*_ with ⟨*η*_*i*_(*t*)⟩ = 0 and correlation *C*_*dyn*_(*t*) ≡ *C*(*t*) − *C*(*t* − *t*′ → ∞) (which goes to 0 at long times).

Time-averaging the log rate of change 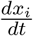 for the persistent types gives results that are independent of Θ with

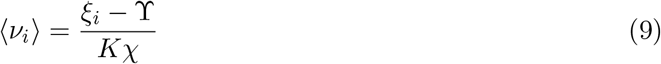

where 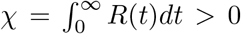 and one readily finds ϒ(*t*) = 0, as expected. Hence only types with positive bias *ξ*_*i*_ > 0 survive. Self-consistency for the total *N* and 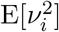 fixes *χ* and the variance of *ξ* [28].

At high temperatures (but still Θ ≪ *K*), averages like ⟨*ν*_*i*_⟩ and ⟨*ν*_*i*_(*t*)*ν*_*i*_(*t*′)⟩ are dominated by rare times at which frequencies are large. Concretely, from Equation 5 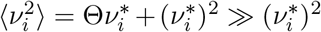. This is dominated by the peaks where *ν*_*i*_ reach the ceiling of ~ Θ/*K* — which occurs a type-dependent fraction of order 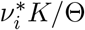 of the time.

For convenience, we temporarily drop the *i* subscripts: characterizing each type by its *ξ* = *ξ*_*i*_, and rescale to 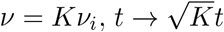 and the other quantities similarly to get rid of factors of *K*. The shape of the peaks in *ν*(*t*), and the ceiling they reach, is caused by the competing effects of a positive instantaneous growth rate 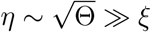 (from ⟨*η*^2^⟩ = *C*_*dyn*_(0) ~ Θ) and the negative feedback from ∫ *R*(*t*−*t*′)*ν*(*t*′)*dt*′. With a peak at time 0, approximating *R*(*t*) ≈ *R*(*t* → 0) = 1 (directly computable) and, crudely, *η*(*t*) approximately constant during the peak, yields *ν* ≈ 2*η*^2^*e*^*ηt*^/(1 + *e*^*ηt*^)^2^ which first grows exponentially at rate *η* (unaffected by *R*), peaks at *η*^2^/2 ~ Θ, turns around in a short *peaking time* 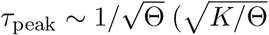 in unrescaled units), and then decreases exponentially. The turnaround is caused by the negative feedback from the other types that do well against it and happen to already be quite abundant, illustrated in Figure 1. A peak occurs whenever *x* is within order one distance from the ceiling. The *η*^2^ height of the peak, together with the Gaussian distribution of *η* causes the upper end of the distribution of *ν* to fall off exponentially as in Equation 5. On long time scales, *x*(*t*) is roughly uniformly spread out over a range of Θ/*ξ*, the fraction of time it is close to the cutoff is the inverse of this.

The typical time, *τ*_eq_, for *x*(*t*) to fluctuate over its full range — loosely the *equilibration time* — is much longer than *τ*_peak_. The behavior on intermediate time scales *τ*_peak_ ≪ t ≪ *τ*_eq_, will be most important — and general. When *x* is far from the upper cutoff, we conjecture (and show in Appendix B.3) that the effects of the feedback will not be large. For intermediate *x*(*t*), the effects of *ξ* will be small as well, and *x*(*t*) will undergo stochastic motion driven by *η*(*t*). The behavior of *C*(*t* − *t*′) on intermediate time scales is dominated by the probability of peaking again at *t*′ after having peaked at *t*: this corresponds to a return process for the random walk in log-space. Self-consistency (Appendix B.2) shows that a conventional random walk cannot be correct and suggests that the typical distance 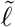 traveled during the walk over a (short) time *t* goes as

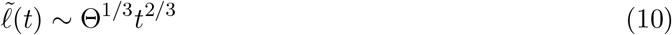

which corresponds to a super-diffusive random walk. The correlation function in the intermediate regime is concomitantly *C*(*t*) ~ (*t*/Θ)^−2/3^, which approaches the right value for *t* ↘ *τ*_peak_. This scaling is confirmed numerically in Figure 7.

The bias will only matter after a timescale *τ*_eq_ ~ Θ ≫ *τ*_peak_ (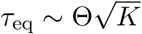 in unrescaled units). On this time scale, the net effects of *ξ*, on average Δ*x* ~ *ξτ*_eq_, will be comparable to the effects of the stochastic *η*(*t*). Over time scales *τ*_eq_, *x*(*t*) will thus typically go from a series of peaks down over its full range to ~ −Θ/*ξ*. Concomitantly, *C*(*t*) will decay over the time scale *τ*_eq_ to close to its long time limit of order unity — a consistency check on the ansatz for 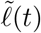. On longer time scales, the balance between the effects of the upward bias and the stochastic *η*(*t*) causes the exponential distribution of −*x* (power law of *n*) in Equation 5. For times longer than *τ*_eq_, the types with anomalously small *ξ* will continue to fluctuate, but now roughly diffusively, 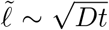 out to 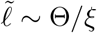 with the diffusion coefficient 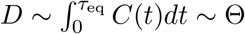.

Similar heuristic arguments yield understanding of the response *R*(*t* − *t*′): its behavior is determined by the effect that a “kick” from *η* at *t*′ has on the probability of a peak at *t*. In Appendix B.3 we show that

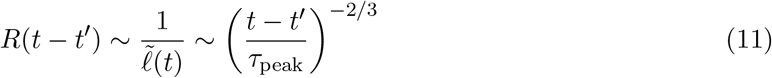

for *τ*_peak_ ≪ *t* ≪ *τ*_eq_. This is order unity at *t* ~ *τ*_peak_ — justifying the approximation that *R*(*t*− *t*′) ≈ *R*(0) = 1 during a burst. But the integral of *R*(*t*) is dominated by *t* ~ *τ*_eq_: the above form yields 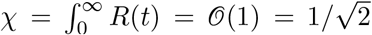 (the actual result from static calculation). In Appendix B.3 we show that, in spite of the slowly-decaying memory, the cumulative effects of the response are small compared to those of the noise, *η*(*t*), except during and soon after peaks. These inferences about the response function validate the consistency of the assumptions and approximations. Knowledge of the form of the snapshot distributions of the frequencies, {*ν_i_*}, has provided powerful hints about the dynamics.

#### 3.2.1 Dynamics for near-antisymmetric interactions

When γ > −1, the stable high-diversity chaos of the perfectly antisymmetric model collapses. The fixed point is generically unstable, and this extends to global instability. The roots of this instability can be understood in terms of the behavior of the purely antisymmetric case. When some type *i* rises to large frequency, it remains large until types *j* driven upwards by positive *V*_*ji*_ themselves rise enough to drive type *i* down via their negative *V*_*ij*_. The rate of this process is set by the strength of negative feedback, proportional to −γ, as well as the typical (log) distance the frequencies need to change by in order to peak. For −γ < 1, he negative feedback is weaker and the bursts are higher and longer. Longer bursts mean that the instantaneous growth rates (set by the highly abundant types) change more slowly as well. All types will thus explore a larger range of abundances — more problematically, reaching lower log abundances. But then it takes longer for types to again burst; this positive feedback loop continues and increases fluctuations — eventually driving extinctions. A quantitative analysis can be carried out when γ = −1 + *ϵ*, *ϵ* small (Appendix B.4), by defining an approximate “energy” *E* in terms of effective 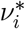 defined by running averages of the *ν*_*i*_(*t*), and by showing that *E* is generically increasing (Figure 8).

In parallel work, Roy et. al. [81] have studied the case γ = 0 and *Q* < *Q*_*c*_ also finding diverging chaos driving extinctions. We conjecture that stable high-diversity chaos is not possible anywhere in the *Q* − γ parameter space except for the special point γ = −1, *Q* = 0.

## 4 Chaotic coexistence stabilized by migration

We have argued that ecological chaos will generally cause larger and larger fluctuations to low abundance, quickly leading to a cascade of extinctions. In order to prevent extinctions an influx of new individuals of each type is needed. The simplest model (used to describe island biogeography [82]) is to consider a “mainland” with a pool of each type from which there is a low rate of migration to the “island” being studied. If the migration is not too small, the dynamics are approximately deterministic and chaos on the island will be stabilized. But this begs the question of how the mainland diversity is stabilized: What prevents global extinctions?

We will show that large collections of identical islands with migration between them can lead to a stable high-diversity spatio-temporally chaotic state with some types going globally extinct, but a large fraction stable for exponentially long times. We develop understanding of this “phase” by building on our analysis of the chaotic phase of the pure ASM, which is special but nonetheless contains key elements of more generalized dynamics.

Consider a large set of *I* islands (or “demes”) with migration between all pairs of them. The islands are identical (types interacting via the same matrix *V* on each) and without spatial structure, with equal migration rates *m* per individual out of each island. (To make clear the role of extinctions, we here work with abundances — population sizes — rather than frequencies). The dynamics of the abundance, *n*_*i,α*_(*t*), on island *α* follows Equation 2 (in terms of abundances) with an additional term for the net migration of individuals, 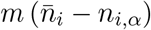 where 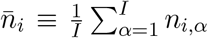 is the average over all islands of type *I*. The Lagrange multiplier, ϒ_*α*_(*t*), keeps the total population, Σ_*i*_ *n*_*i,α*_ = *N*, the same on each island separately.

With large *m*, the population dynamics on the islands would synchronize and behave like a single island with diverging chaos driving global extinctions. But as long as the migration rate is lower than the largest Lyapunov exponent for chaos on one island, the chaos on each island will be desynchronized [83]: we will show that this leads to global stability. We are interested in migration rates much smaller than rates of changes of populations on an island. A characteristic scale for the exponential growth or decay rates of local populations is the rms magnitude of the time-averaged interaction Σ_*j*_ *V*_*ij*_ ⟨*n*_*j,α*_⟩/*N*, which we denote 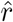, of order 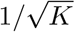 for order one interactions.

Since the total number of migrants into island *α* is proportional to the island-average 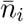 and not the instantaneous population on that island, *n*_*i,α*_, even with small *m* the inflow of migrants can compensate for a negative local growth rate to keep the local population at or above a minimum size, 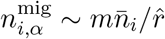, which we call the *migration floor*. We make the ansatz (and establish its self-consistency) that for globally-persistent types, the migration influx, 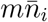, is much larger than 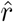, so that *n*^mig^ is still many individuals while much smaller than the overall average population per type per island, 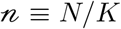. Population fluctuations are then negligible and we can treat the dynamics as deterministic. But this breaks down if 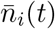 becomes anomalously small: this breakdown will set the dynamics of global extinction.

Simulations with a modest number of islands and types already indicate much of the behavior of the spatio-temporally chaotic phase. Some types go extinct globally (even in the deterministic approximation) with the total population of those types, 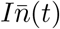, decreasing roughly exponentially with time as shown in Figure 2(a). But a majority of types persist for long times, even when there are local extinctions. The dynamics of the subpopulations of a single persistent type across all the islands is shown in Figure 2(b). While much of the time the population on an island is near the migration floor as indicated, on each island there are *local blooms* up to high abundance which produce enough emigrants to other islands to avoid local extinctions and enable later blooms on other islands. Indeed, even a small initial population on just a single island can, if it arrives at an anomalously favorable time, bloom and seed other islands leading to long term persistence, as in Figure 2(c). It is only when these blooms are too rare that global extinctions occur. Therefore the statistics and dynamics of the blooms control the behavior of this phase.

**Figure 2:**
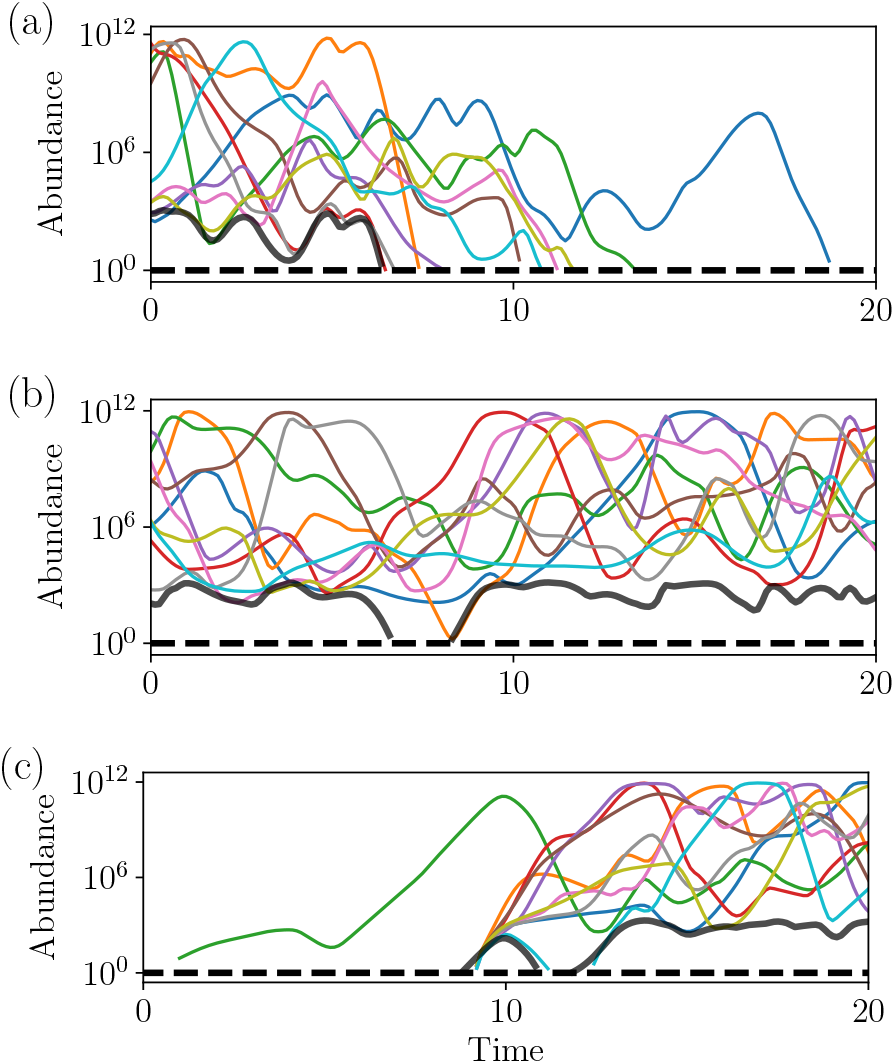
*Abundance trajectories across islands for* single *types* across *I* = 10 islands with *K* = 75 initial types (γ = −0.8, *N* = 10^12^, 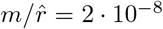). Black lines show the *migration floor* at which the local populations are stabilized by migration from other islands, and the dashed line the boundary for extinctions. (a): For this type, a global extinction results from a cascade of local extinctions due a shortage of large blooms on enough islands. (b): A persistent type in steady state. Local extinctions can occur when the migration floor, and hence some local populations, drops below *n* = 1. But migration prevents local extinctions from becoming a global extinction as long as a bloom to high abundance is occurring on at least one island. (c): An invasion can start from a small local population on one island that, if lucky, initiates a bloom from which migrants establish populations on other islands, and leads to long-term persistence.

An important property of the spatio-temporal chaotic phase is the distribution of abundances. A snapshot of the abundances on a single island shows that these are distributed approximately uniformly in log(*n*) over a range down to the migration floor, with additional weight at the low end. This is seen from simulations in Figure 3, together with a comparison to the ASM behavior. But the average abundance of each type over all islands 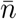, or averaged over long times on one island, will be dominated by the blooms to high abundance. Like in the ASM, the average populations of different types vary much less than the abundances in a snapshot.

**Figure 3:**
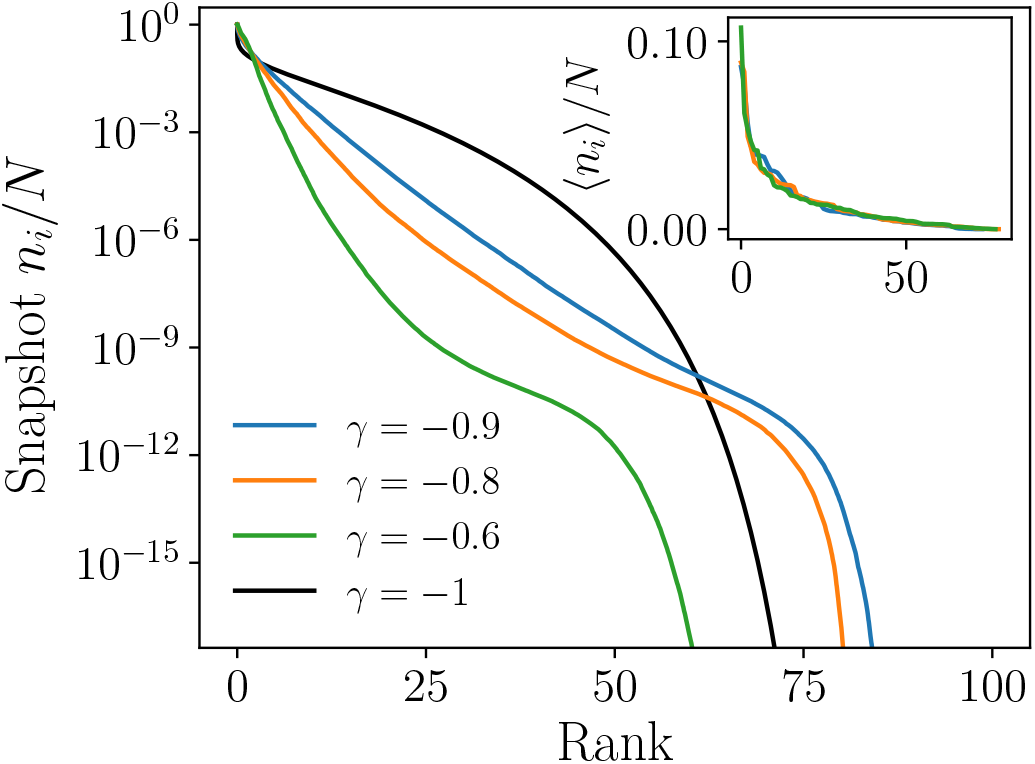
*Rank-frequency plots for island model* (*K* = 100 types, *I* = 50 islands, migration rate 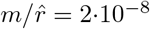, and various γ). “Snapshots” on a single island show fractional-abundances broadly distributed over log space with some clustering around 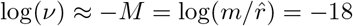 due to stabilization via migration. Sharp fall-off for very low abundances is due to types on their way to extinction. Time or island averaged abundances show much narrower distributions, (inset, on linear scale) with little γ dependence. For comparison, a snapshot of the single-island ASM with Θ = *M* and 85 persistent types (black) has similar intermediate abundance but different behavior at high and low abundance.

A striking feature of the spatio-temporally chaotic phase is that much of the behavior is essentially independent of the migration rate as long as it is very small. The time scale for the blooms is proportional to log 1/*m*, and the lower limit of the local populations is proportional to *m*. But the statistical dynamics over a wide range of log(*n*) and the distributions of the island averaged populations, 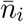, depend only weakly on even log(*m*). The number of types, *K* sets the overall time and population scales, but otherwise plays little role as long as it is sufficiently large.

We will obtain overall understanding of this phase by analyzing the behavior in the limit of an infinite number of islands and infinite population size on each, assuming a steady state, and showing that this is consistent. Then we will analyze the behavior for large but finite *N* and *I*.

### 4.1 Analysis of spatio-temporally chaotic phase

With the chaos on each island out of sync, in steady state each island should behave similarly, and averaging over very many islands is equivalent to averaging over the chaos on one of the islands.

We can make a dynamical mean-field approximation for a single type, *I*, on one island, *α*, as in Equation 6 with the additional migration term and an additional self-consistency condition that the time average on each island is equal to the island average. To find a steady-state, one assumes an island average 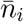, does the local analysis to find the time average on one island, ⟨*n*_*i,α*_⟩ as a function of 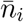, and then find the solution of 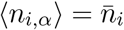. There are two possibilities in steady state: globally extinct types with 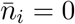, and persistent types with 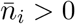. Which occurs will depend on the average growth or decay rate of the population of type *I* under the effect of interactions with all the others on the same island: i.e. the bias *ξ*_*i*_, which is the same on each island.

The net bias on type *i* is 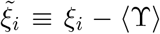 with the Lagrange multiplier ϒ(*t*) being positive (to counter the weaker feedback at high abundance than for −γ < 1). Since the biases {*ξ*_*i*_} have zero mean, less than half the types have positive 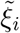. Nevertheless, we shall see that many of the types with negative net bias survive: this is the key to the stabilization of diversity by spatiotemporal chaos. We focus on γ strongly negative, but not too close to −1 (for which crossover effects complicate matters).

The behavior of persistent types is qualitatively similar to that of the purely ASM, but with the crucial difference of stabilization at low abundance by migration. Types with negative net bias spend a fraction of the time on each island near to the migration floor, with occasional local blooms to high abundance. The migration to other islands is dominated by these blooms, and is otherwise negligible: the statistics of the blooms is thus key. The crucial feature, mathematically, is that the log-abundances *x*(*t*) = log *n*(*t*) are broadly distributed, so averages are dominated by the large fluctuations and ⟨*n*(*t*)⟩ = ⟨*e*^*x*(*t*)^) ≫ *e*^⟨*x*(*t*)⟩^.

At the height of a bloom, a type can have multiple peaks to high abundance, reaching a size 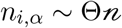, where Θ is loosely analogous to the “temperature” in the ASM. Abundances of typical types with net bias not far from zero will be spread out roughly uniformly on the logarithmic scale from the peaks down to the migration floor, thus over a range 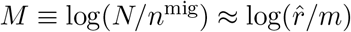. The parameter *M*, which we are taking to be large, determines many of the quantitative properties of the chaotic phase. From the uniform distribution, peaks would occur a fraction of time of order 1/*M*: for consistency, these must have the right height implying that Θ ~ *M*; i.e. the “temperature” is set by the migration rate. With very many persistent types, a considerable number will be at large abundance on an island at any time, and the mean-field approximation should be good. Note that in our simulations, even for quite large *K* only a few types may be abundant at each time, but averaging over the duration of the blooms, during which many types will peak, is roughly consistent with the mean-field results, Thus capturing the properties of a type by its 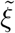, is a reasonable approximation.

The correlation and response functions are dominated by the statistics of the peaks, qualitatively similar to the single island ASM. The typical duration of individual peaks is 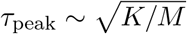. Away from peaks *x*(*t*) undergoes a super diffusive random walk, with exponent 2/3, on time scales up to 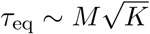 over which it ranges down to the migration floor. The rough balance between the stochastic fluctuations and the average effects of the net bias suggests that unless the net bias, 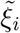, is strongly negative, blooms will not be rare and will typically last times of order *τ*_eq_.

As already illustrated in Figure 3, the snapshot of the abundances on one island are broadly distributed, while the average over all islands 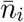 varies much less and is of order 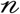 with a coefficient determined by 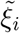. For positive 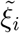, the migration plays little role, and the behavior is similar to the ASM with 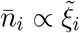. For negative bias, 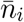 can be much lower but still be non-zero, as analyzed below. Although the bias of a type is not directly measurable one can infer the 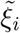 using the mean-field structure (below). Simulations exhibit the expected features of 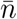 as a function of 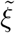, shown in Figure 4.

**Figure 4:**
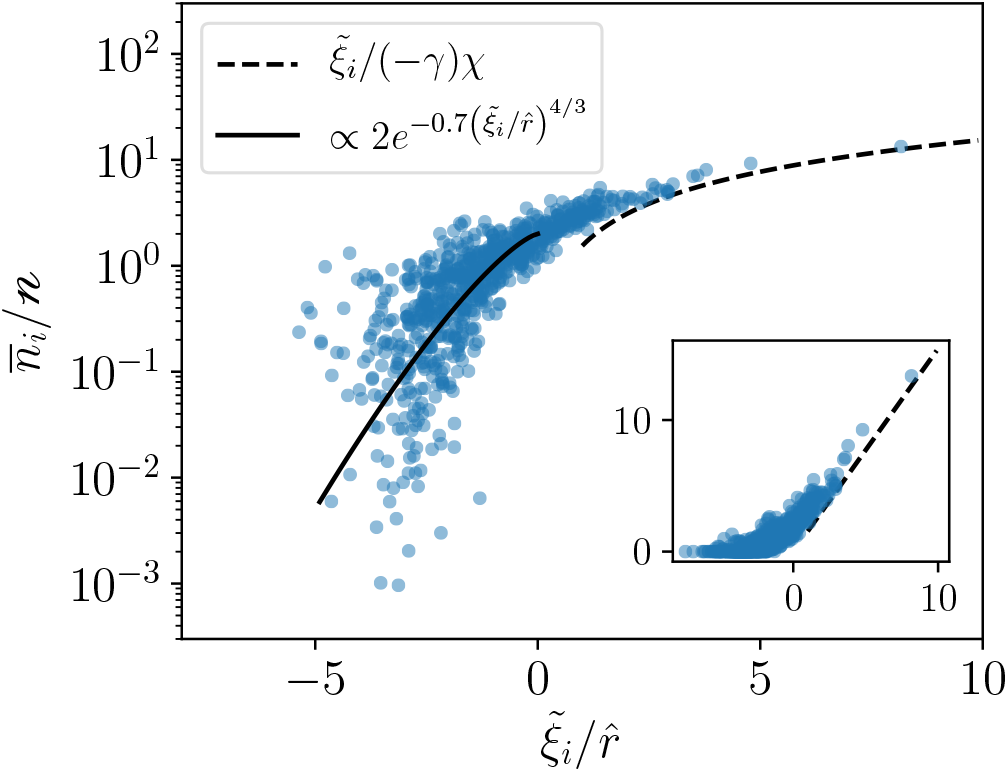
*Island-average abundances vs. net bias*. Mean-field analysis predicts the relationship between abundances (log scale) and net bias 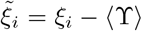 (average growth/decay rate of type *i* at low abundance). Inset: same plot in linear scale. Biases estimated from 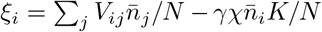 and normalized by the typical growth (or decay) rate, 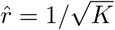, with abundances normalized by average across types, 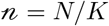. Theoretical predictions for *ξ*_*i*_ substantially positive (dotted line) and substantially negative (solid line) agree reasonably well with the numerics. Data shown for the long-term persistent types for γ = −0.8, *K* = 100, *M* = 18 and *I* = 24 islands. The dependence of 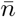 on *ξ* is largely independent of *I*, although fewer types survive for smaller *I*.

#### 4.1.1 Persistence and averages with infinitely many islands

We now investigate the conditions for extinction and persistence and determine the form of 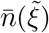. The island-averages are most readily understood quantitatively for types with either much larger than typical net biases (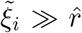, the standard deviation of the {*ξ*_*i*_}) or large negative net biases, 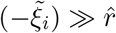. For large positive biases, the effects of migration are small and we expect the behavior of *n*_*i,α*_(*t*) to be similar to the ASM with Θ ~ *M*. The migration term is negligible unless 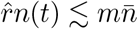, which occurs with exponentially small probability 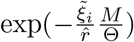 similar to the ASM. In terms of the mean-field quantities, the island-average is 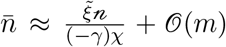 as in Equation 9, which, for positive net biases, compares well to 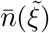 in Figure 4. From simulations the biases are inferred from the time-averaged interactions, Σ*V*_*ij*_ ⟨*n*_*j*_⟩ /*N*, equivalent to 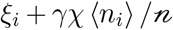 in the mean-field.

The susceptibility *χ* is chosen such that the median of the inferred *ξ*_*i*_ is zero, as expected with the extinct types included.

For large negative biases, 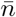 is determined by the probability of rare blooms to large abundance during which the stochastic fluctuations of *η*_*i,α*_(*t*) must overcome the negative bias. For a bloom resulting in a large log-abundance change Δ*x*, large deviation analysis is carried out in Appendix C.1. This is analogous to the probability of a random walk going “uphill” in a potential, but with the noise here being power-law temporally correlated, as in Equation 10. We find that the bloom probability for 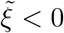 is

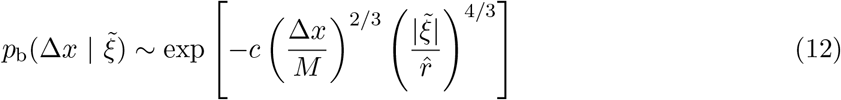

with the *c* (here and henceforth) an order unity coefficient. The start-to-peak time of the least unlikely bloom is 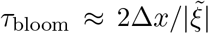 of order *τ*_eq_ for typical negative 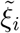. The important blooms start at the migration floor at 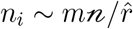 and increase to a maximum of 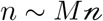 (limited by the response of the other types) a change of Δ*x* ≈ *M*. The island average abundance is proportional to the probability of such maximal blooms: 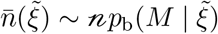, dropping powers of 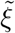 and other factors that depend on more detailed dynamics during the blooms, including the multiplicity of peaks for each bloom, as in Figure 1. The plots of 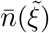 in Figure 4 show the predicted roughly exponential dependence on 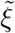 when it is negative. But we see that, since Δ*x* ≈ *M*, for typical negative 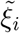, the exponential factor is not particularly small. Thus the types with negative net bias will contribute substantially to the total population. Since *M* drops out of the statistics of the blooms, much of the behavior is only very weakly dependent on the migration rate — especially as *M* itself depends only logarithmically on *m*.

For very negative 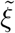, the probability of blooms, and hence 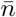, decreases enough that the migration floor will be much lower than typical. Then even on a log-scale, the blooms will have to go up by substantially more than *M*. Eventually, for negative enough 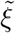, the duration of the blooms becomes of order *τ*_eq_. Since the power law correlations of the noise are cutoff for times greater than *τ*_eq_, (and the dynamics becomes like diffusion with a bias) the bloom probability drops more steeply with Δ*x* and a self-consistent solution for 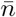 cannot be found. This occurs for 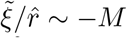 with extremely small 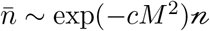. Thus, in principal only a fraction of order 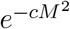 of the types go globally extinct with both *N* and *I* infinite (numerically, less than 1% for the parameters used in Figure 4). But in practice, the least abundant persistent types will easily be driven extinct by either finite population-size, or finite number-of-islands, fluctuations as analyzed next. Nevertheless, this analysis does predict that with low migration, a substantial majority of the types can be globally stabilized, most with 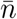 a substantial fraction of 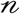 but a tail extending down to smaller 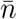 even on a log-scale, as seen in Figure 4.

#### 4.1.2 Persistence with a finite number of islands

With a finite number, *I*, of islands, the average across islands, 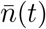, will fluctuate, as seen in the abundance trajectories of a single type across 10 islands in Figure 2. With many islands, the blooms occur often enough to maintain 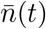 roughly at its infinite island value, 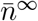. To maintain 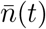 and prevent global extinction, at least one island must typically be blooming at a time. For a type with large negative bias, a single bloom starting from the migration floor must increase by at least Δ*x* > *M* to maintain the same 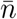. At any time, the probability of such a bloom occurring on at least one island is roughly 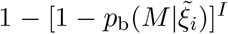. Thus for 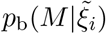 less than some constant times 1/*I* — corresponding to 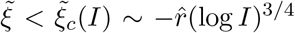 — blooms cannot sustain the population which will hence fluctuate downwards to global extinction.

But types with somewhat negative 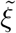 will still be stabilized by migration: this can already be seen with two islands in a diffusive approximation for the effects of the other types. Consider a new type invading at low enough abundances that the response term in Equation 6 due to other types is negligible. The coupled dynamics of a single type across two islands can be analyzed exactly, as carried out in Appendix C.2. The growth rate of 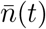 when invading is positive for 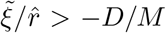 where 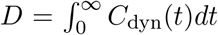 is the diffusion constant, of order 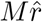. Already with two islands, 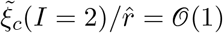 suggesting that a majority of types are stabilized. These results assume a stable temperature for the chaotic dynamics, but in practice more than two islands are typically needed to prevent diverging fluctuations. However fewer than ten islands are sufficient to stabilize a majority of types across the range of γ and *m* studied.

### 4.2 Stochastic global extinctions

We turn now to the problem of survival with a large but finite population *N* on each island. Fluctuations in 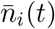 allow local extinctions to occur (for islands with low and decreasing abundance at that time) when the migration floor 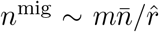 becomes of order one individual. As shown in Figure 2, a bloom on another island can later drive *n*^mig^(*t*) back to being large and re-establish the local population. But if no blooms occur for an extended period of time, local extinctions will happen on islands that are not yet repopulated when the local conditions would have caused a bloom: this can lead to a runaway to global extinction. We need to understand the probability of this occurring as a function of the number of islands, the net bias of the type of interest, and a key parameter: the typical migration floor log-abundance, 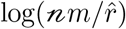.

With infinite or a very large number of islands, the averages across islands do not fluctuate much so the condition for global survival is simply that *n*^mig^ ≫ 1. For *Nm* very large, analysis of the bloom probability in Equation 12, shows that all but a fraction of order 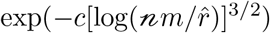 will survive. We see that this results in a peak in the fraction of persistent types at small, but not extremely small, *m* (Figure 5).

**Figure 5:**
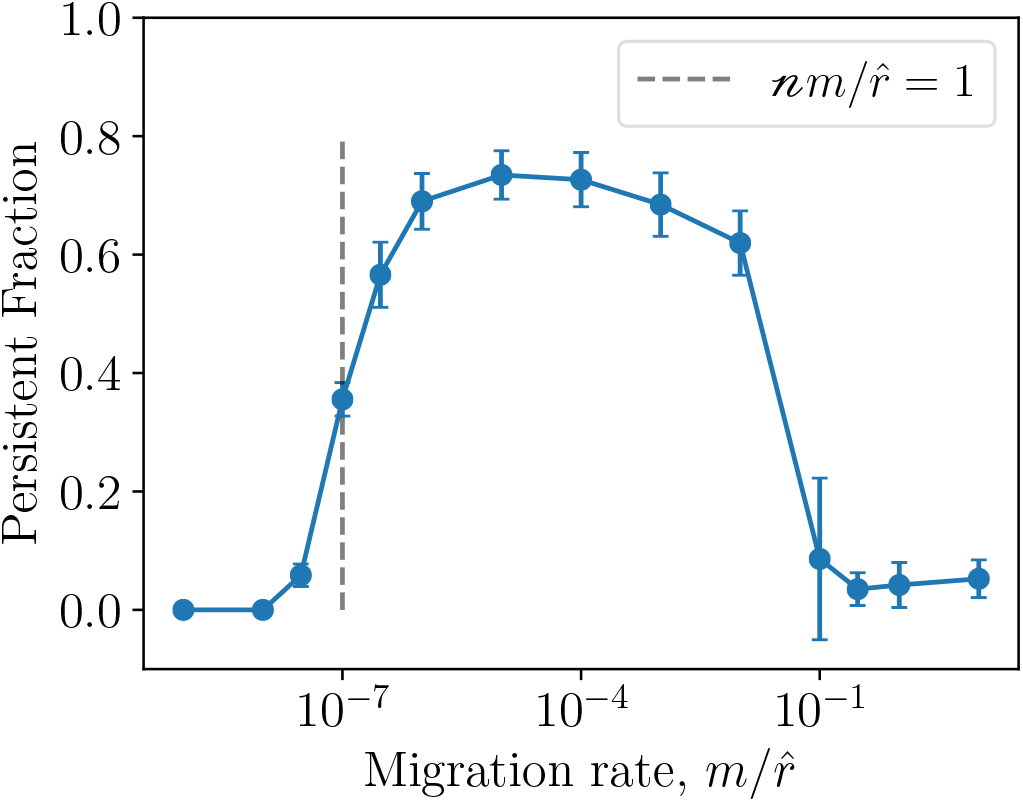
*Persistent fraction of types vs. migration rate*. Out of *K* = 100 types the persistent fraction is greatest for intermediate migration rate. For 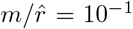, islands become synchronized leading to a large loss of diversity, while for 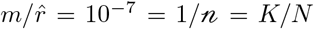 the migration floor for a typical type drops below one individual. Results averaged over 20 instantiations of *V*_*ij*_ with population per island, *N* = 10^9^, and *I* = 30 islands. To avoid issues with stochastic extinctions at finite times, simulations were run without an extinction threshold and then the persistent types defined as those with a migration floor, 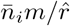, greater than one. This is exact in the limit of many islands.

With a moderate number of islands, stochastic extinction can occur if, by bad luck, the migration floor fluctuates down to *n*^mig^ ~ 1 for long enough that abundances on all islands fail to reach large size for an extended time. The dynamics of the migration floor will consist of periods of downwards drift and recovery by blooms. Since the crucial blooms that reach high enough to set *n*^mig^ typically start from the migration floor, the dynamics of *n*^mig^(*t*) depend on its value at earlier times, complicating the analysis. We make a conjecture for the least-unlikely trajectory of 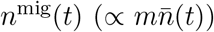 to global extinction and evaluate its probability: we expect that this will yield an estimate of the correct form. A natural choice is to assume that the least unlikely trajectory is log *n*^mig^(*t*) decreasing steadily at some rate *v*: this would be correct if its dynamics were driven by Gaussian noise plus an average upwards bias (like the log-populations on single islands). Typically 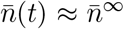, so a fluctuation of 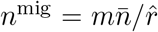 down to 1 would take a time 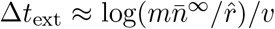 to extinction.

The extinction probability is dominated by the likelihood that in the time Δ*t*_ext_, no island has a bloom large enough that it would make *n*^mig^(*t*) increase above its steady decline, which would require the log-abundance of the bloom, Δ*x* = log *n*(*t*) − log *n*^mig^(*t*), to be at least *M*. In the frame of the decreasing log *n*^mig^(*t*), the bloom has to overcome a less negative effective bias, 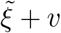. Since blooms on different islands, or at well-separated times, are roughly independent, the probability of extinction due to having no blooms for a time Δ*t*_ext_(*v*) is roughly

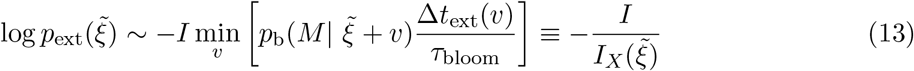

where the factor Δ*t*_ext_/*τ*_bloom_ is roughly the number of possible blooms on each island. For large negative bias 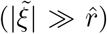, the least-unlikely speed is 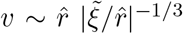, small enough that the bloom probability, *p*_b_, is little-changed from its *v* = 0 value and we find that the characteristic number of islands is

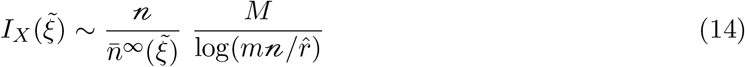

(up to unknown powers of 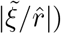). The second factor, coming from Δ*t*_ext_/*τ*_bloom_, is the ratio of log(*N/n*^mig^) to log(*n*^mig^/1), so it compares the log-drop from *N* to the migration floor to that from the migration floor to the extinction threshold.

Stable types will survive for exponentially long times proportional to 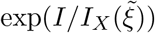, while marginal types will go extinct much more quickly. This can be seen by plotting the fraction of initial types that have gone extinct versus time (Figure 6); even with a modest number of islands many types survive for long periods of time. The exponential dependence on *I* can be seem explicitly by plotting the *k*th extinction time as a function of *I* (inset).

**Figure 6:**
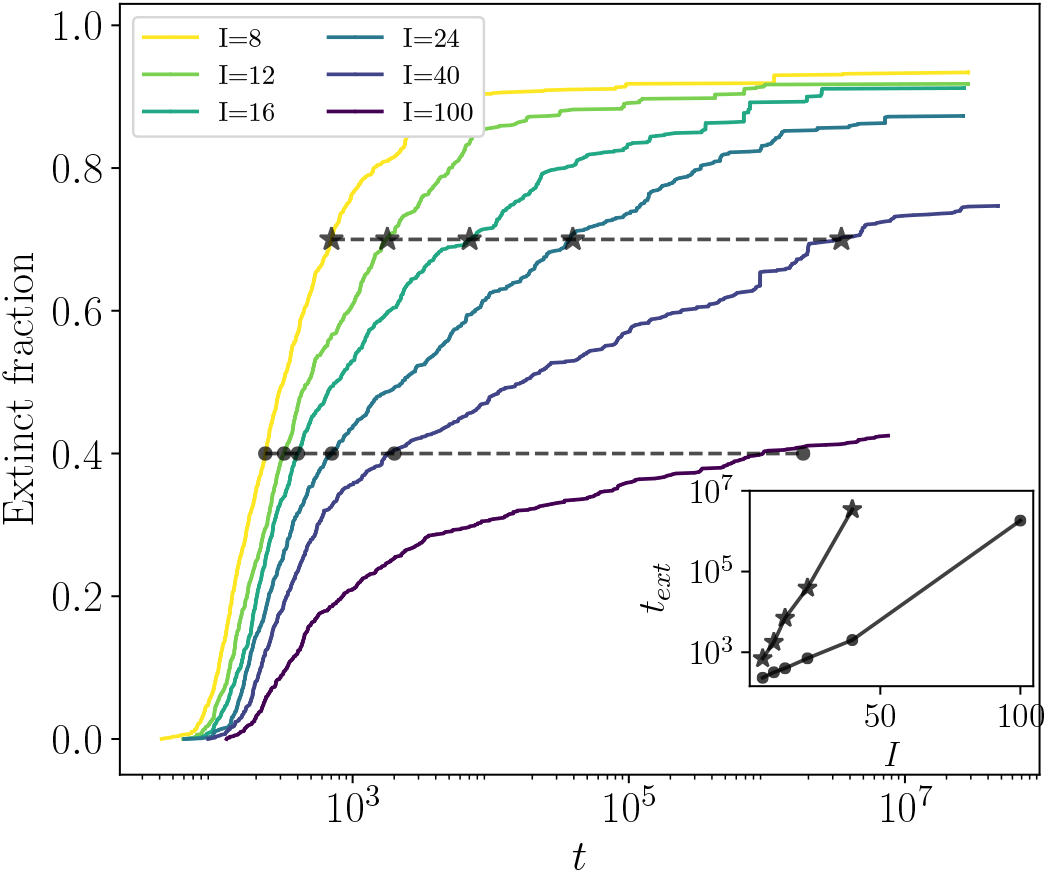
*Extinct fraction vs. time:* After rapid initial extinctions, remaining types persist for long periods of time. With increasing number of islands, *I*, more types survive for long periods and the typical time taken before a fixed fraction of types have gone extinct (horizontal lines for fractions 0.4 and 0.7) is exponential in *I* (inset). Data from long simulations of 10 replicates with *K* = 100 starting types, 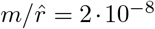 (*M* = 18), *N* = 10^13^, and γ = −0.8.

## 5 Discussion

We have shown that antisymmetric correlations in the Lotka-Volterra interaction matrix, together with simple spatial structure, are sufficient to stabilize extensive diversity of an assembled community. No niche-like assumptions or special properties are needed. This spatio-temporally chaotic “phase” is very robust; the key ingredient is the negative feedback induced by the antisymmetric correlations and sufficient — albeit very small — migration. While some fraction of the types go deterministically extinct, a majority, including types.with substantially negative average growth rate, persist for very long times. The key phenomenon underlying this stability is that each type occasionally has a local bloom to high abundance which provides migrants to the other islands. As these blooms are nearly independent from island-to-island, global extinctions occur only if blooms do not happen on any of the islands for a sufficiently long period of time. This yields persistence for times exponentially long in the number of islands.

### Generalizations

Much of our analysis has focused on the asymptotic but unrealistic regime when logarithmic functions of the population size and the migration rate are large. This has enabled us to obtain many results in a general framework based on dynamical mean-field theory. Yet our simulations show that the predicted behaviors are correct, even semi-quantitatively, for modest sizes of parameters. A crucial prediction is the exponential scaling of survival times with the number of islands, illustrated in Figure 6 for realistic parameters: total population per island, *N* ≈ 10^13^ — of order the number of human gut microbes [84] — *K* = 100 types, and the ratio of the migration rate, *m*, to the typical local population growth (or decay) rate, 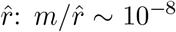 (corresponding to the logarithmic parameters 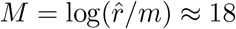, and 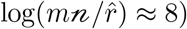.

The diverse spatio-temporally chaotic phase should exist far beyond the models we have analyzed. In Appendix D we discuss the behavior of the general random Lotka-Volterra model with niche-like interactions (parameterized by *Q* = −*V*_*ii*_) and argue that the spatio-temporally chaotic phase exists over much of the γ − *Q* phase diagram. With selective differences *s*_*i*_, the number of types coexisting is limited when 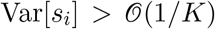 (Appendix D.3), but the behavior otherwise similar. More generally, the phase should exist with sparse interactions, broad distributions of interaction strengths, correlations due to phylogenetic relatedness of types, or with some variations between islands. Indeed, even overall antisymmetric correlations of the interactions are not essential. The key is the absence of types that always outcompete most other types, and that for each type there are some ecological interactions that provide negative feedback preventing it from persisting at high abundance. We showed this effect via antisymmetric structure of the interactions (Section 3), loosely inspired by host-pathogen dynamics, but this can also occur with moderately strong niche-like competition with its own type 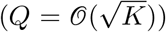 combined with sufficiently random interactions with many other types. Indeed, in parallel work, Roy et. al. [81, 85] have found, for γ = 0, a chaotic phase stabilized by migration with behavior consistent with our predictions.

Dynamics with large swings of local abundances are crucial, but these need not be driven primarily by complex ecology. If environmental fluctuations that temporally benefit some types more than others occur on all islands roughly independently but statistically similarly, then blooms on different islands will be roughly independent and limited by the overall carrying capacity on each island. A low rate of migration can then stabilize the coexistence of diverse strains even if they all do badly on average: the time-averaged fitness plays the role of the net bias, 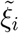 which can be negative for most of the long-term persistent types. This is in striking contrast to the behavior of a well-mixed population in a fluctuating environment whose dynamics is controlled by the exponential of the average fitness. For the island model, instead, the global population dynamics of the spatio-temporal ecosystem is controlled by the average (across islands or time) of the exponential of the fitness (which is much larger). Thus the basic picture the spatio-temporally chaotic phase and its striking consequences for diversity will be very general.

### Abundance distributions

A key quantitative characteristic of the spatio-temporal chaos is a general consequence of the dramatic fluctuations of local populations. “Snapshots” of the local abundances will be broadly distributed on a logarithmic scale (Figure 3). How universal are such abundance distributions? At low abundance, they will be affected by details of migration, while at high abundance, different mechanisms for limiting blooms (Appendix D), such as strong niche-like interactions (Appendix D.2), or some types doing atypically well on average (Appendix D.3), will change the abundance distribution. It is the wide intermediate range of abundances that will be far more universal – and this is given by the dynamics of the seemingly-special perfectly antisymmetric model! The lack of universality at both high and low abundance, means that summary statistics like the “Shannon diversity” (entropy of the distribution) or “species richness” (total number of types observed) are poor characterizations of the abundance distributions: the former is dominated by high abundance types and the latter by very low abundance types.

The more universal intermediate part of the abundance distribution will be approximately a power-law with slope one. This is the same as from the neutral theory of ecology [33, 38, 86]. In fact, a neutral model with immigration from a mainland at a rate 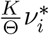 has an identical joint distribution of frequencies as the ASM with temperature Θ and fixed point frequencies, 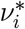. The underlying dynamics, however, are very different, qualitatively and quantitatively, especially for large microbial populations for which birth-death fluctuations are usually negligible. Many eco-systems exhibit broad abundance distributions which are used to argue for/against different ecological scenarios [15, 36]. But our work surely implies that sampling at one location and time is far from sufficient: spatio-temporal data — especially by time series from deep sequencing — are needed to distinguish between different scenarios.

### Bacteria-phage diversity

The most natural context for antisymmetric correlations in the interactions between types, is diverse strains of both a host species and a pathogen species, especially bacteria and phage. Models of such systems with perfectly antisymmetric (γ = −1) interactions have been studied previously but with only tens of types [72, 73, 87, 88]. Dynamical mean-field analysis enables understanding of large numbers of types. The primary additional parameter is the ratio of time-scales over which the differences between types result in substantial abundance changes for the bacterial strains vs for the phage strains. If this ratio is unity, the mean-field dynamics of both the bacterial and phage strains is *identical* to the ASM (as shown for fixed points in [27]), and if the time scales differ substantially, we expect quantitative, but not qualitative, differences. Spatial structure obviates the need for perfect antisymmetry — unrealistic in any case — and our basic picture of the spatio-temporally chaotic phase should hold.

Dynamical diversity from bacteria-phage interactions has recently been studied by other approaches. For example, some explicit “kill-the-winner” models treat phage predation as stochastic events leading to bacterial population collapse [70, 89]. While these exhibit persistent diversity, it is unclear whether there is a reasonable underlying population dynamics that could give rise to the caricature used. The advantage of models with explicit population dynamics for bacteria *and* phage, is that avoidance of extinctions cannot be put in “by hand”. In addition, models of the type we studied naturally allow for (and indeed, may require) non-specific interactions, instead of the one-bacteria-one-phage scenario of the original kill-the-winner models [59, 60, 90], which destabilizes in the presence of demographic stochasticity. Indeed, non-specific interactions may be needed to stabilize a chaotic phase [69].

### Extensions

Our theoretical and computational frameworks should enable biologically-relevant extensions in various directions.

### Realistic spatial structure

The effects of real spatial structure surely merit exploration. With conditions being the same everywhere, but transport either by local diffusion or occasional long-range “jumps” such as by wind, ocean currents, or hitchhiking on animals [91], the process of recovering from local extinctions is more complex as it must involve spatial propagation of repopulation “fronts”. Can a substantial fraction of many similar types survive globally even in a system of infinite spatial extent?

### Microbial spores or “seedbanks”

When faced with unfavorable environmental conditions, many microbes enter a reversible state of dormancy by forming spores or other resting structures [92]. This “microbial seedbank” of dormant cells has been estimated to make up 80% of cells in soil and 40% of cells in marine environments [92] and is suggested to be an important component in the persistence of rare bacteria [14]. The formation of long-lived — but not immortal — spores is another mechanism that can stabilize a diverse ecologically chaotic phase, indeed, even in a well-mixed system with no spatial structure. If some fraction of each type form spores that die off slowly but occasionally germinate to produce actively dividing cells, the spore population effectively averages the chaotic dynamics of the active cell populations over time, in analogous way to the set of islands. The product of of the average spore lifetime, *τ*_*sp*_, and the typical growth or decay rates, 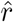, of the active cell populations, is roughly equivalent to the number of islands. With deterministic dynamics and statistics of interactions similar to those we have studied, a substantial fraction of the types will survive forever. Even with spore extinctions the survival times will be exponentially long in the the spore-lifetime, including for types whose net bias — average growth rate of the active population, 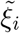,— is negative. Individual types will persist for time 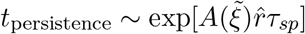 with *A* depending also on the population size of the spores, the germination rate, and other factors. Thus even spores with moderate lifetime can yield a diverse chaotic ecology with many types that have negative average growth rate surviving for extremely long times.

### Phenotype models

An unsatisfactory feature of Lotka-Volterra models of many interacting types is that the organisms do not interact via their phenotypic properties: their phenotype is defined by their interactions with others. There is a long history of more explicit phenotype modeling for species interacting via consumption of common resources — MacArthur consumer resource models and generalizations [20, 21, 22, 23, 24, 65]. More generally, one can consider interactions via many chemicals in the environment with the dynamics of the populations set by the chemical concentrations, and the dynamics of the chemical concentrations set by the population sizes of the various types. Our analysis should be readily extendable to at-least-simple-versions of such models, most simply when they reduce to Lotka Volterra models with interactions determined by the effect of one type on the environment, and the response of the other type to this. Under what conditions will a spatio-temporally chaotic phase exist in ecological models with interactions only via modifications of the environment? And how much diversity can be stabilized by the interplay between chaos and migration? This is a productive avenue for future research. A more general question is how systems will behave with direct interaction between pairs of individuals — and thus again Lotka- Volterra structure — but with the interactions determined by phenotypic properties of the two organisms.

#### Evolution and ecology

The most important feature left entirely out of this work is the evolutionary dynamics: here (and in most other work on random Lotka-Volterra models and generalizations) we have assumed that there exists a large number of closely related types and asked what happens when a community of these is assembled. But can such a state evolve by a “natural” evolutionary process in a broad class of models? Although in reality for microbial populations there is not usually a clear separation between ecological, migrational, and evolutionary time scales, and others have shown how diversity can be established and maintained in pathogen-host models with very rapid evolution [69]. But observationally, coexisting strains of the same species are found to have a wide spectrum of genetic differences and hence times since their common ancestors. Thus the most basic question is what happens when the evolution is the slowest process: this is the hardest case for extensive diversity. Models should characterize the types by phenotypic properties which are what evolves, and the interactions be determined by these — without strict tradeoffs, as is often assumed [23, 93]. Rather, there might be a low likelihood of generalist mutations (higher *s*_*i*_) in an already-evolved population, with a much higher likelihood of mutations that take advantage of the particular combination of types in the system at the time and location at which they arise.

Can a highly diverse chaotic community evolve in such models? If so, will the system undergo continual Red Queen evolution with no systematic “improvement”? Or will the evolution tend to get slower and slower? The statistical properties conditioned on evolution will likely be quite different than those in an assembled community. What general features might emerge? And how might different scenarios — and different possible “phases” of the eco-eco dynamics — potentially be distinguished by data from real microbial populations?

## Acknowledgments

AA was supported by a Bowes BioX fellowship and Stanford’s Center for Computational, Evolutionary and Human Genomics, MTP by William R. and Sara Hart Kimball as a Stanford Graduate Fellow and by a National Science Foundation Graduate Research Fellowship, DGE-114747, and all authors by the National Science Foundation via PHY-1607606. We thank Giulio Biroli and Felix Roy for valuable discussions and sharing the results of their parallel work. Computer resources were provided by the Stanford Research Computing Center’s Sherlock cluster.

## A Numerics

Simulations were carried out using the standard Runge-Kutta method (explicit RK4) applied to the basic dynamical equations in Equation 2 and its extension to the island model with a step size small compared to the smallest timescale, *τ*_peak_. Times *t* in all plots correspond to this scaling, except where otherwise noted.

For the simulations with islands, local extinctions and re-establishments due to new migrants from other islands were treated deterministically: a good approximation when the growth-decay rate varies greatly. Extinction occurs when the abundance of a type drops below one: *n*_*iα*_ < 1. Re-establishment occurs when the influx of migrants, 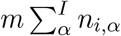, exceeds the basic rate of change 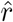 (equal to 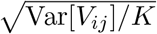) for the population dynamics. This condition corresponds to the migration floor, 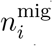, becoming greater than one. In our regime of interest when the typical migration floor, 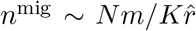, is large compared to one, the deterministic approximation changes some quantitative details but not the key results.

## B Single-island mean-field theory

## B.1 Mean-field rescaling

For the calculations in the rest of this section (as well as some in the main text), it is useful to rescale the variables so that the factors of *K* vanish, and so that the mean-field equations are in terms of the abundances *n*_*i*_ instead of the frequencies *ν*_*i*_. We define the dynamics as

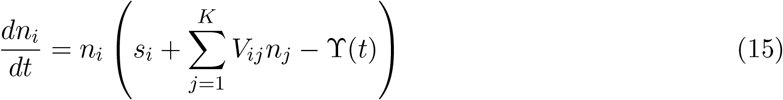

and make the choice 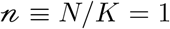. In order to get growth rates of 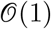, we set Var[*V*_*ij*_] = 1/*K*.

This gives us the mean field equation

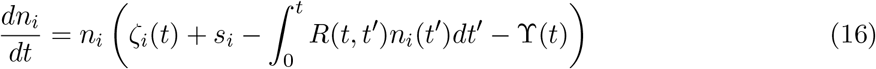

where

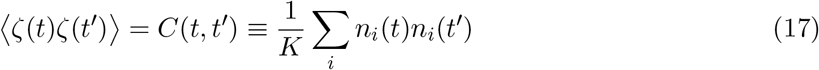

and

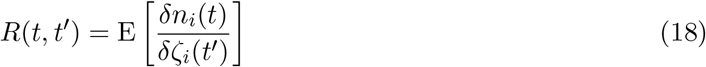

This obviates the need to carry around factors of the typical growth/decay scale 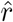 or, equivalently, the factors of *K*.

## B.2 Super-diffusive correlation function

The mean-field dynamics of the ASM (Equation 6) requires self-consistently solving for the correlation function *C*(*t* − *t*′). In general there is no analytical closed-form solution, but progress on understanding the scaling behavior of *C* (and the response function *R*) can be made by making an ansatz for the dynamics and checking for self-consistency.

The very short (*t* = *t*′) and very long (*t* − *t*′ → ∞) time behaviors of *C*(*t* − *t*′) are determined by the static distributions (Equation 5). The intermediate time behavior of *C*(*t*−*t*′) can be determined by approximating the dynamics of the log-abundances *x*(*t*) as a random walk in the appropriate regime. If the noise correlation *C*_*dyn*_(*t* − *t*′) were integrable and equal to a diffusion coefficient 2*D*, then the typical scale of *x*(*t*) − *x*(*t*′) would be 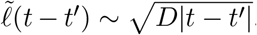. But *C*(*t*, *t*′) ~ Θ^2^Prob[*n*(*t*) ~ Θ & *n*(*t*′) ~ Θ], i.e. that *n* is at a peak at both times. If *x*(*t*) were undergoing roughly a random walk with an upper boundary (at which it “bounces off” due to the effects of *R* analyzed in B.3), this probability would decay as 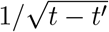. But then 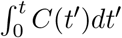 would grow rapidly with *t* so the diffusive ansatz is inconsistent.

We instead conjecture that for 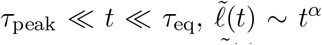 with 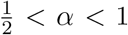. A time *t* after a burst, *x*(*t*) will be spread roughly uniformly down to 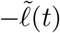 so has probability of order 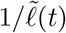 of again being near the cutoff and hence in another burst. This yields 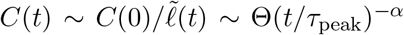 (analogously to the diffusive case). But ⟨(*x*(*t*) − *x*(*t*′))^2^⟩ is given by the double integral of the correlations of *η*(*t*) which are ∫ ∫ *C* ~ (*t* − *t*′)^2−*α*^. Thus self-consistency requires *α* = 2/3 so that *x*(*t*) undergoes fractal Brownian motion with

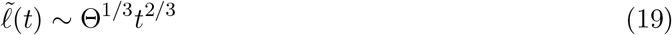

which is, as it should be, of order unity for *t* ~ *τ*_peak_ ~ Θ^−1/2^. The correlation function in the intermediate regime is thus *C*(*t*) ~ (*t*/Θ)^−2/3^. A time *τ*_eq_ after a peak, *x*(*t*) will have fluctuated down to of order −Θ. At this point the intermediate-time scaling regime breaks down and *C*(*t*) is of order unity — as it must be to match the long-time behavior 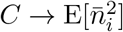.

The scaling for intermediate times can be confirmed numerically (Figure 7) by computing *C*(*t*) − *C*(*∞*). After the initial sharp drop on the short time scale, *τ*_peak_, the correlation function drops off roughly as *t*^−2/3^ for a time of order *τ*_eq_, as expected.

**Figure 7:**
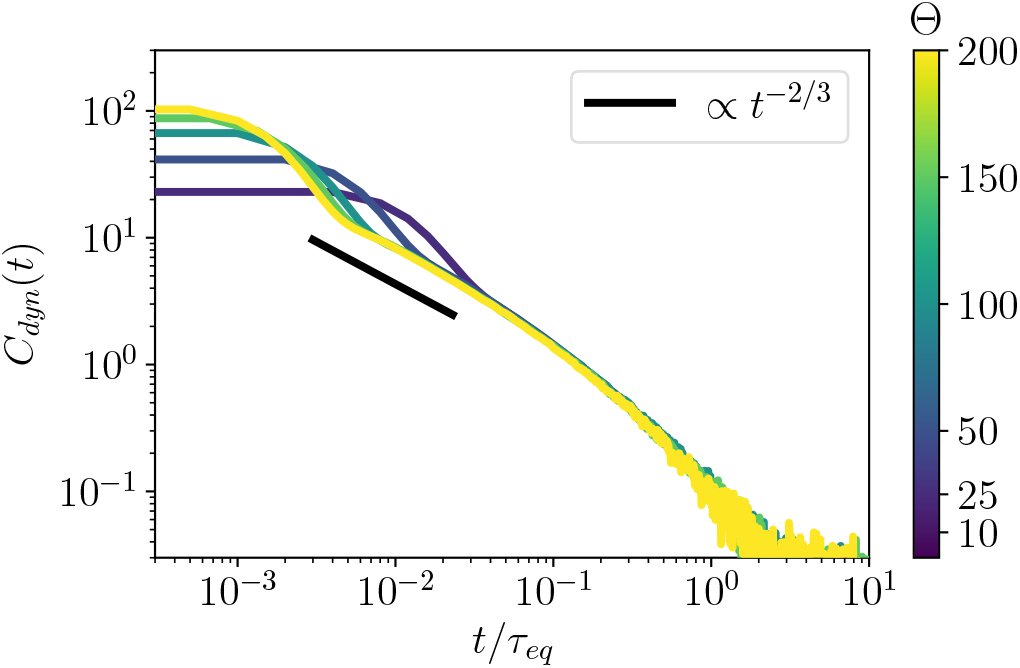
*Correlation function for the antisymmetric model (ASM)* with *K* = 201 persistent types. Rescaling by the long timescale 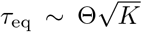 collapses the curves of *C*_dyn_(*t*) for different “temperatures”, Θ. At intermediate times, the correlation function decays roughly as a power law of *t* consistent with the predicted *t*^−2/3^, from Equation 10. The short time correlations decay faster for higher temperatures as expected from the scaling of the timescale of a peak, 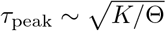.

## B.3 Scaling form of the response function

The response function for the ASM can be understood quantitatively using similar heuristic reasoning to the analysis for the correlation function. If at time zero *x*(0) = −*y* is far from either cutoff, a small kick upwards from a delta-function spike of *η* will have little effect on *n*(*t*) until a time at which 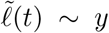 after which the distribution of *x*(*t*) will be spread out over a range of width 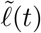 from the ceiling down. If it happens to be at a peak (*n*(*t*) within a factor of order one of the ceiling) this will contribute of order the peak height, Θ, to *R*(*t*). Ignoring memory of prior history, the average contribution to *R* from such an initial condition is roughly 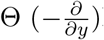 Prob[near peak at *t* given *x*(*t* = 0) = −*y*] which can be approximately integrated over the distribution of *y* to yield

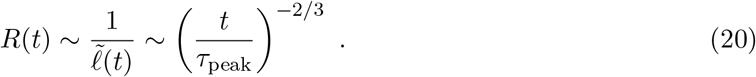

For *t* of order 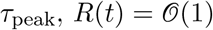 consistent with its *R*(0) limit of exactly 1.

The effects of the long-term memory embodied in *R*(*t*) can now be estimated. After an initial peak at time zero before which *x*(*t*) had not been small for some time, we can crudely estimate 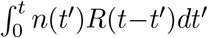 from an average of the multi-peak *n*(*t*′), ⟨*n*(*t*′⟩ ~ Θ (*t*′/*τ*_peak_)^−2/3^ (as in derivation of the correlation function in Appendix B.2) and *R*(*t* − *t*′) from Equation 20 yielding ∫ *Rn* ~ (*t*/Θ)^−1/3^ which is larger than the upwards bias until *t* ~ *τ*_eq_. But at shorter times, the cumulative effects of the − ∫ *Rn* term in *dx*/*dt*, is Δ*x*_*feedback*_(*t*) ~ (*t*/*τ*_peak_)^2/3^ — the same order as the cumulative effects of the stochastic driving, 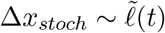. Thus we expect that the approximation we have made of ignoring the feedback except in the peaks, should give the correct scaling behaviors. A crucial self-consistency check is that the integral of *R* over times up to *τ*_peak_ (beyond which it decays more rapidly), is of order unity: the exact result being 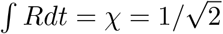. This consistency could not have occurred for any different scaling of 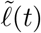: thus we could have obtained the key result 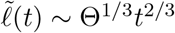 directly by matching the short and long time limits.

## B.4 Diverging chaos with close-to-antisymmetric interactions

For γ = −1 + *ϵ*, *ϵ* small, the dynamics of the Lotka-Volterra model (Equation 2) is similar to the pure antisymmetric (γ = −1) model with some temperature Θ on short timescales, determined by the initial conditions. However, on long timescales, as we will show, this temperature increases.

For convenience we work with the dynamical equations in terms of the abundances *n*_*i*_, with *N* = *K* and Var[*V*_*ij*_] = 1/*K* so that the relevant timescale for the γ > −1 behavior is 1/*ϵ*.

Consider a system initialized with an equilibrium distribution (corresponding to γ = −1) at some initial temperature Θ_0_. We make the ansatz that there is some adiabatic timescale *τ*_*ϵ*_ ≫ *τ*_eq_(Θ_0_) ~ Θ_0_ over which the distributions of types do not change much. We will check this assumption for self-consistency later. We define the adiabatic average abundance 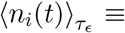 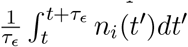, as the time average over this adiabatic timescale.

We can define an effective energy *Ẽ* by replacing the 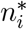 by the adiabatic averages 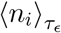:

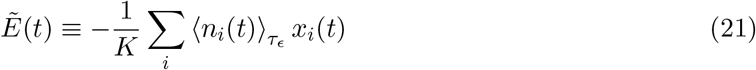

where *x*_*i*_ ≡ ln(*n*_*i*_), and we anticipate the (slow) time dependence of *Ẽ*. We conjecture that the joint distribution of the *n*_*i*_ (really, the *x*_*i*_) is close to that of the equilibrium distribution for the γ = −1 case with Θ ~ *Ẽ*(*t*) (to be checked later). Under this assumption, computing the dynamics of *Ẽ* will let us show that the “temperature” of the ensemble is increasing.

The time derivative of the energy, 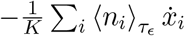, evaluates to

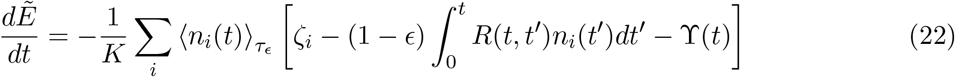

From mean-field identities, we have

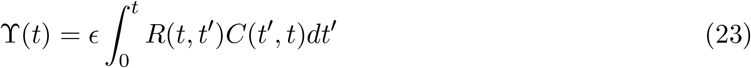

Averaging over types, and taking advantage of the fact that the *n*_*i*_(*t*) sum to *K*, we have

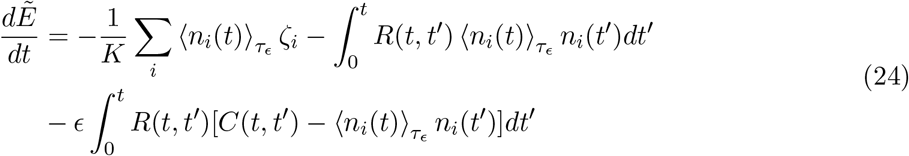

We average *dẼ*/*dt* over *τ*_*ϵ*_. Consider the first term. For the antisymmetric model, the (infinite) time average is

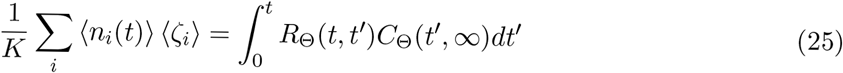

where *C*_Θ_ and *R*_Θ_ are the correlation and response functions for the antisymmetric model at temperature Θ. Since *τ*_*ϵ*_ ≫ Θ we have 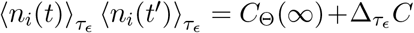 for some small correction 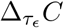 corresponding to the change in *C* from *C*_Θ_ (whose magnitude will be calculated later).

Therefore, we will write:

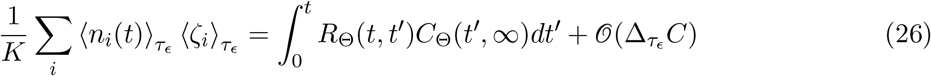

where we conjecture that all the replacements accrue error of the same order 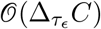.

Similarly, we replace 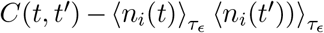 with *C*_*dyn*,Θ_(*t*) = *C*_Θ_(*t*) − *C*_Θ_(∞), the dynamical correlation function for the antisymmetric model. We then have

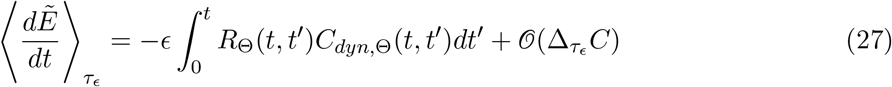

The integral is order 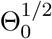 at initialization. If we assume that the current *Ẽ* corresponds to the temperature Θ of the closest antisymmetric distribution, then we have

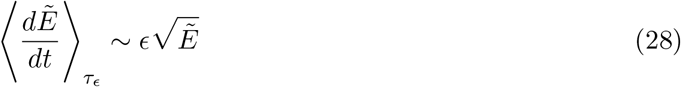

which suggests *Ẽ* ≈ *Ẽ*_0_ + *aϵ*^2^*t*^2^, which can be confirmed numerically (Figure 8).

**Figure 8:**
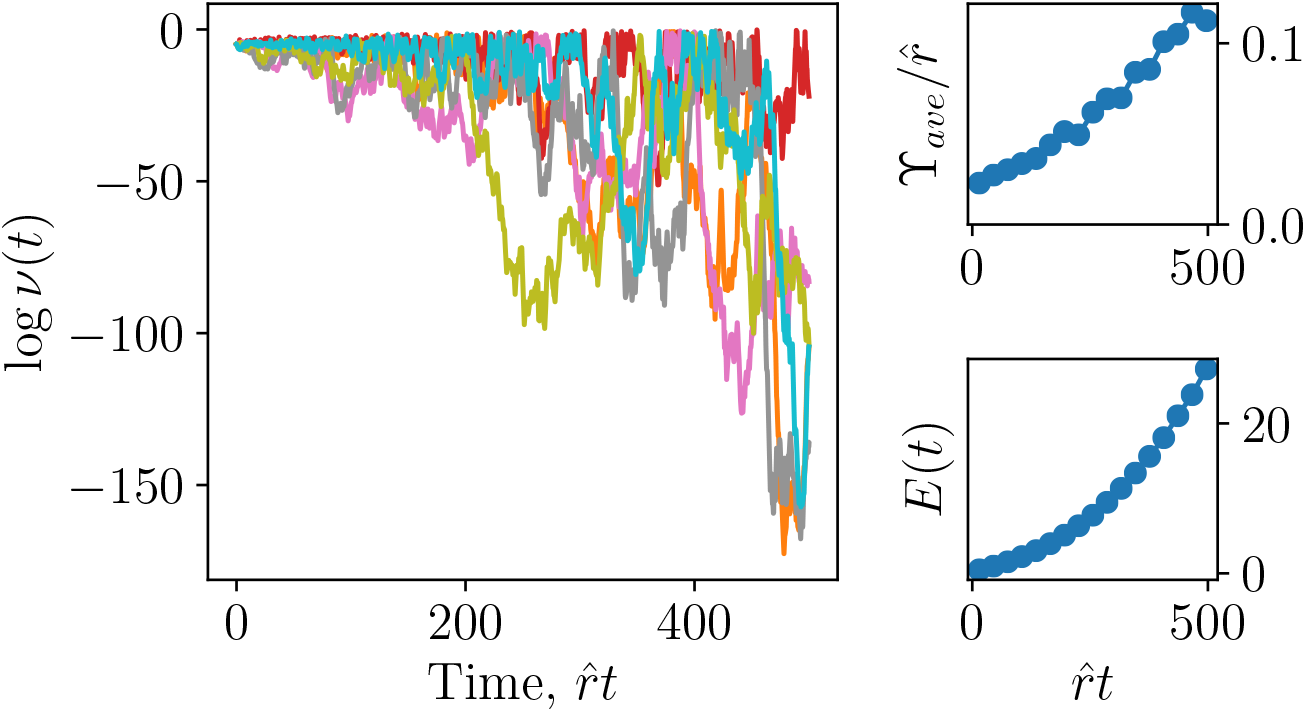
*Unstable chaotic dynamics on an isolated island* (without migration) for a nearly antisymmetric interaction matrix, (γ = −0.99). Starting from the fixed point 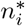 of the antisymmetric part of the interaction matrix, (*V* − *V*^*T*^)/2, the abundance swings become steadily wilder on a log-scale (left, for single simulation). The Lagrange multiplier averaged over the ensemble, ϒ_ave_(*t*), grows roughly linearly (top right) and the “energy” 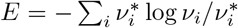 grows roughly quadratically in time (bottom right), as predicted by Equation 28. Simulations with *K* = 301 initial types and averages over 50 instantiations. Time plotted in units of basic growth rate 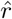.

We can now check that there truly exists an adiabatic timescale *τ*_*ϵ*_, such that 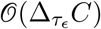 is small. To begin, we compute 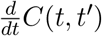. We have

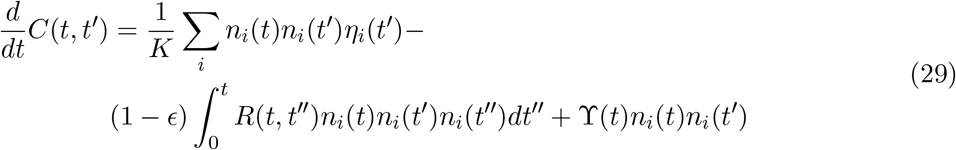

Define 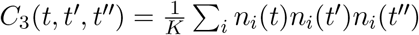. By using mean-field identities, we haves

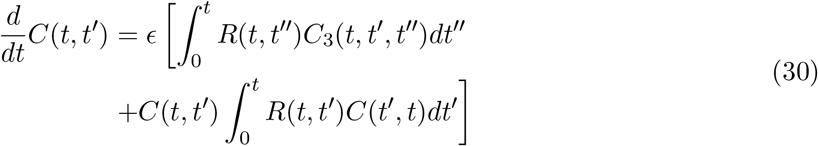

Note that this expression is, so far, exact in the mean-field.

For times of order *τ*_*ϵ*_, we can evaluate Equation 30 using the values of *R*, *C*, and *C*_3_ from the γ = −1 case with temperature Θ. Note that *C*_3_ peaks at Θ^2^ if all three times are within Θ^−1/2^, and at Θ if only two times are close. We have, in a scaling sense,

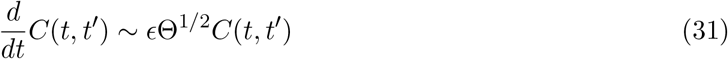

We are now in a position to check the self-consistency of the adiabatic approximation. The adiabatic timescale exists if there is only a small fractional change in *C* over *τ*_*ϵ*_, which suggests *τ*_*ϵ*_ ≪ *ϵ*^−1^Θ^−1/2^. We also require that *τ*_*ϵ*_ ≫ Θ, the timescale for the antisymmetric system to reach steady state. This means that *τ*_*ϵ*_ exists as long as *ϵ* ≪ Θ^−3/2^. For small enough *ϵ*, the adiabatic regime holds and the temperature is increasing.

By the above argument, the temperature increase is guaranteed only until *Ẽ* changes by 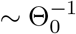 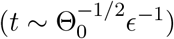. However, the numerics suggest that the scaling form holds for times at least *O*(*ϵ*^−1^) (so that Δ*Ẽ* ~ 1). *Ẽ* continues to grow beyond that (Figure 8), albeit without the *t*^2^ form. With a finite population size, this leads to extinctions and low diversity.

## C Island model mean-field theory

## C.1 Self-consistency analysis for determining island averages

In this appendix we self-consistently find the average abundance across islands, which in steady state will not fluctuate in the limit of infinite number of islands. To do this for types with negative net bias, we need to analyze their bloom probability.

For large negative biases, the probability of a bloom can be estimated from a large deviation analysis of the random walk dynamics of *x*(*t*) ≡ log *n*(*t*), except that the driving noise is power-law correlated. Consider a bloom resulting in some large change in log-abundance, Δ*x*. Except perhaps at the peak, the abundance is very small and the response feedback negligible. The most likely trajectories from the migration floor to a peak near the ceiling will not have intervening returns to the migration floor. And they will have last-peaked long enough ago for the effects of feedback to be negligible. Thus we can consider the large Δ*x* tail of a process driven solely by the net bias plus dynamic noise, 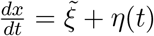, with 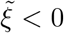 < 0. Since the noise is Gaussian the distribution of Δ*x* after a time Δ*t* will also be Gaussian

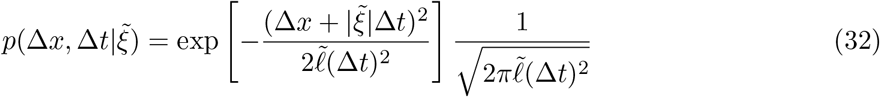

where 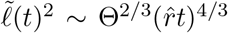 is the variance of 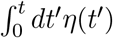 in the power law regime of *C*_dyn_(*t*), as in Equation 10.

The probability of a bloom of fixed size Δ*x* is the integral over all times, but this is dominated by the most likely star-to-peak time, readily seen to be 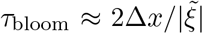. Plugging in, the most likely bloom has probability

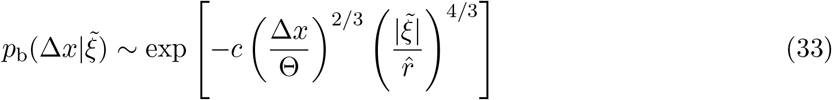

with some order one constant *c* and with non-exponential powers of 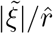 dropped.

Starting from the migration floor, a bloom of size Δ*x* reaches an abundance 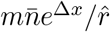. This grows faster with Δ*x* than the bloom probability decays with Δ*x*, so the average is dominated by the largest possible bloom, with Δ*x*\*, that reaches a maximum abundance 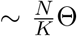 due to the feedback of other types. The average abundance due to the blooms is

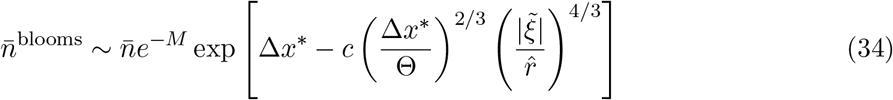

dropping non-exponential factors and using the migration floor, 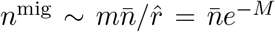. As the average will be dominated by these blooms, self-consistency implies that 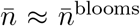 so that the expression in square brackets in Equation 34 must be ≈ *M*. But since Θ ~ *M*, for 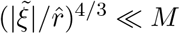 — i.e. the vast majority of negative-bias types — the log-change in a bloom is roughly the same for all these, with

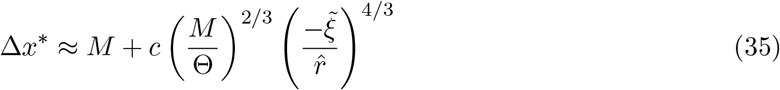

not substantially larger than *M*. Hence we conclude that

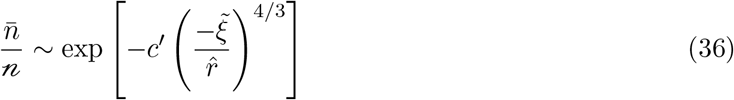

which is only reduced from the type-average abundance, 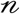, by a substantial factor for atypically negative net biases: 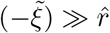.

For the rare types with strongly negative net bias, 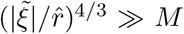, blooms must be substantially larger with 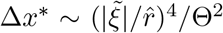. Such blooms take correspondingly longer time, with *τ*_bloom_ becoming of order *τ*_eq_ ~ *M* for biases 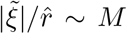. For such long blooms, the noise correlations crossover from the power law regime to the diffusive with 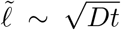 with diffusion coefficient 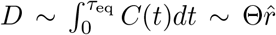. The bloom probability is then simply 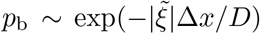. The self-consistency condition for the average reduces to 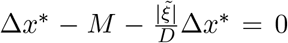, which has no solution for 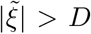 or 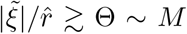. Thus there is no self consistent 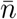 for large negative biases. But the fraction of extinct types is very small, scaling as exp(−*cM*^2^) as inferred from the Gaussian distribution of 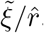.

## C.2 Two island toy model

The stabilization of chaos by migration for a finite number of islands can most easily be shown, semi-quantitatively, by considering just two islands and approximating the dynamical effects of the other types by white-noise. The key is to analyze either invasion or extinction starting from very low abundance of the type of interest in which case the response terms in the mean-field equations are negligible. A simplification then occurs because the difference in log-abundances between the two islands, *y* ≡ *x*_1_ − *x*_2_ = log(*n*_1_/*n*_2_), has dynamics decoupled from the mean of the logs, *z* ≡ (*x*_1_ + *x*_2_)/2. The invasion eigenvalue – mean growth rate when rare — is 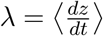, which can be evaluated by averaging over *y* to determine the condition for persistence vs extinction.

The mean-field dynamics for the log-abundance are 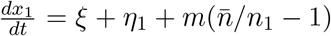 and similarly for *x*_2_. Plugging in 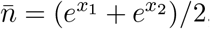, the migration term simplifies to 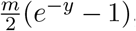, independent of *z*. The dynamics of *y* therefore decouple:

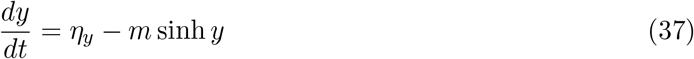

where the noise *η*_*y*_ ≡ *η*_1_ − *η*_2_ has correlation function 2*C*_dyn_(*t*).

In the diffusion approximation for the noise, the Fokker Planck equation can be solved for the stationary distribution,

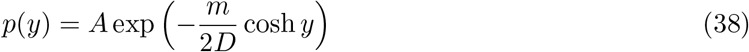

with diffusion constant 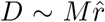. In the asymptotic regime *M* ≫ 1, this distribution is flat until it is cutoff at *m* cosh *y* ~ *D* or *y* ≈ ±*M* (dropping log *D* factors) so the normalization constant is *A* ≈ 1/2*M*. As expected, migration acts to keep the islands’ log-abundances within *M* of each other.

The mean log-abundance, *z* = (*x*_1_ + *x*_2_)/2, has dynamics

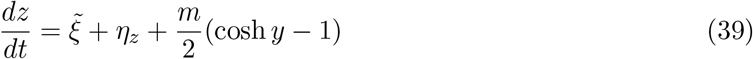

With *η*_*z*_ mean zero and independent of *η*_*y*_. Thus the invasion growth rate is 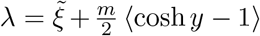. By symmetry, ⟨*e*^*y*^⟩ = ⟨*e*^−*y*^⟩ so ⟨cosh *y*⟩ = ⟨*e*^*y*^⟩ ≫ 1. The average is dominated by the largest values of |*y*| where cosh *y* ≈ *e*^|*y*|^/2 is a good approximation. So from the stationary distribution in Equation 38, the average is

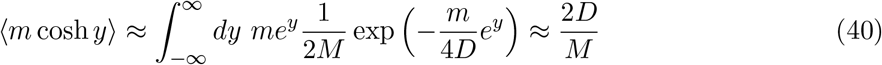

The invasion growth rate is positive if 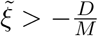 and since 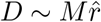, this becomes 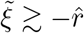. Surprisingly a substantial fraction of the negative net bias types can survive even with only two islands.

## C.3 Persistence of marginally stable types with finite number of islands

For each type with a given 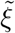 there is critical number of islands 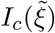 such that the type goes extinct for *I* < *I*_*c*_ even with infinite population size. Conversely, for each *I*, there is a critical bias for persistence: the analysis above (Appendix C.2) shows this for *I* = 2.

In Section 4.1.2 we explained that there needs to be at least of order one island with a bloom at any time to avoid global extinction. Thus the critical number of islands should be roughly 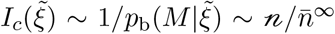 (from Equation 12), where the 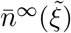 is the average with an infinite number of islands. When *I* is just above 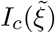, that type has roughly one island blooming at a time and successive blooms occur roughly *τ*_bloom_ apart. For these barely stable types, the island average 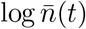 fluctuates by a large amount, of order *M*, between blooms. The next bloom is likely to start from the migration floor when the previous bloom is peaking. This is because the migration floor follows the bloom closely (but shifted down by *M*) when there is only one bloom at a time.

We can approximate such marginally stable dynamics by discrete steps from one bloom to the next. A bloom of size Δ*x* starting from 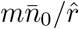, produces an average

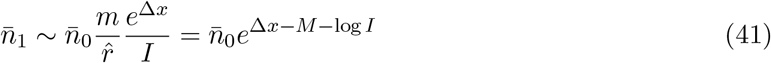

The relevant bloom is the largest one occurring across the *I* islands, with distribution

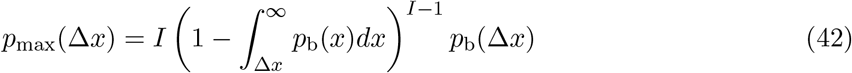

where *p*_b_ is the bloom probability in Equation 12.

The typical size *B* of the largest bloom can be estimated from the condition that the probability of larger blooms, 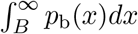, is approximately 1/*I*. This gives

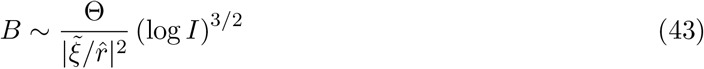

Since the distribution of the largest bloom size falls off rapidly for Δ*x* < *B* and is approximately *p*_b_(Δ*x*) for Δ*x* > *B*, the standard deviation is roughly the decay scale in *p*_b_,

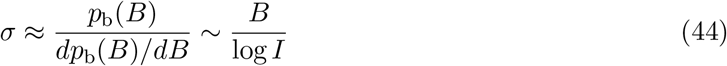

From bloom to bloom, the island average will undergo a random walk with diffusion constant *σ*^2^ and drift-bias *w* ≡ *B* − *M* − log *I*. This drift bias is close to zero because the types are barely stable, so *B* ≈ *M*. If the number of islands is only slightly more than the critical number, 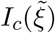, the drift bias is roughly

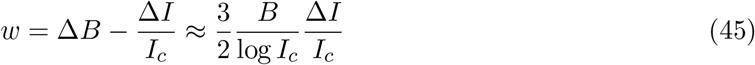

from evaluating the derivate, where 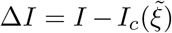. The probability for a random walk fluctuation of the migration floor a log-distance 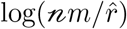 from the typical *n*^mig^ to extinction is

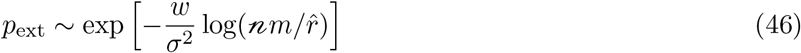

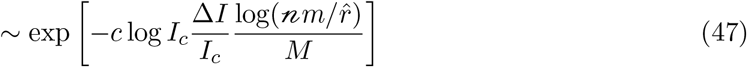

with *c* some order one constant. The extinction probability has a similar form to the result for strongly stable types but is exponential in Δ*I* instead of *I*.

## D Model generalizations

## D.1 Generalizations of random Lotka-Volterra models

In the main text, we restricted analysis to the behavior of random Lotka-Volterra models with strong antisymmetric correlations (γ close to −1), focusing on the case without large self-interactions 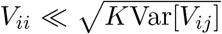 or significant selective differences: 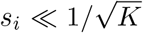. We here discuss consequences for the “phase diagram” of random Lotka-Volterra models more broadly.

A distribution of fitness differences *s*_*i*_ can readily be incorporated into our analyses. Their primary effect is to reduce diversity by *ξ*_*i*_ + *s*_*i*_ now determining ⟨*n*_*i*_⟩. An important po*√*int is that, for fixed total population size, *N*, the effects of random interactions decrease as 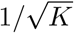 due to averaging over *K* types, while fitness differences are determined by a *K*-independent factor, their standard deviation 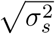. Thus the behavior will depend on 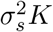. Quantitative analysis of the ASM (Appendix D.3) shows that, for Gaussian-distributed *s*_*i*_, the number of persistent types goes as *K*/2 for 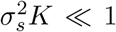, and 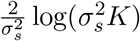 — a much smaller but still potentially large number — for 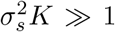. Therefore a large number of types can still survive even with significant *s*_*i*_’s. In the island model, types with substantially negative total biases, 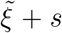, can be stabilized by the migration. An interesting possibility (which we have not yet explored) is that this may make the total population become dominated by the negative bias types such that the persistent fraction in the island model scales differently with 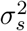 than in the ASM.

We next consider changing the symmetry parameter, γ. As γ approaches 0 from below, simulations show that with increasing probability the diverse chaotic phase collapses to a fixed point or limit cycle with a handful of types surviving, as seen in Figure 9. This collapse can occur when the chaotic dynamics takes the system close to a fixed point or limit cycle which can exist if for some subset of types the symmetric part of their sub-matrix of *V*_*ij*_ produces ϒ large enough to suppress the other types. Even for more negative γ, the symmetric part can lead to large fluctuations in ϒ in the chaotic phase, requiring very large *K* to reach the asymptotic regime of many types simultaneously at large abundance.

**Figure 9:**
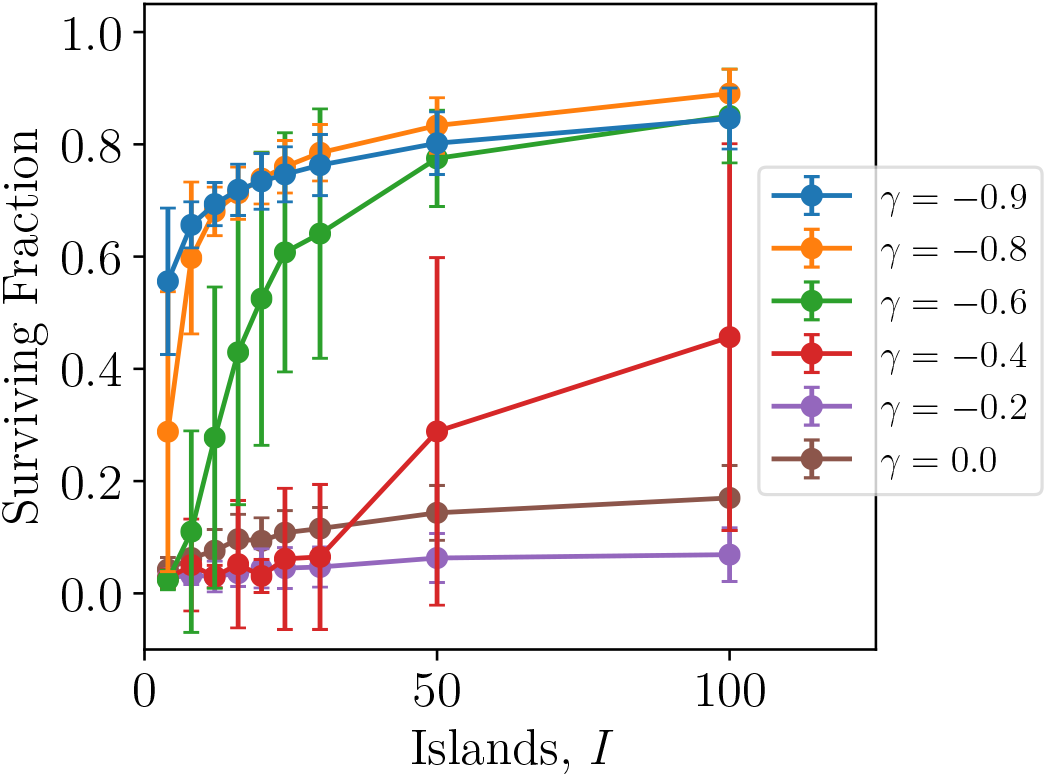
*Fraction of types surviving vs. number of islands*, of *K* = 100 initial types, and a range of symmetry parameter, γ, values. Results averaged over 20 simulations with *M* = 18 (*m* = 2 *·* 10^−8^).

Most recent work using has focused on stable communities caused by niche-like self interactions, 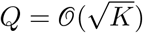 [26, 28, 30, 31, 75]. (Note that [31, 75] use a model with a third parameter but this can essentially be eliminated by use of the Lagrange multiplier, ϒ(*t*).) For 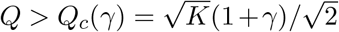 there is a unique large stable community, with 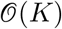 types surviving. In the island model, this will be the same on each island. The absence of persistent dynamics clearly distinguishes this phase from our chaotic phase. But even without dynamical information, the phases are very different: the abundance distributions in the large stable community are much narrower — truncated Gaussian — rather than being broad on a logarithmic scale.

## D.2 *Q* − γ phase diagram

What happens in the rest of the *Q* − γ phase diagram? Rather than studying the full island model, it is easier to study a mainland model with fixed mainland populations of equal size 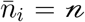 that prevent all extinctions. Preliminary indications from our simulations are that a diverse chaotic will exist for all *Q* < *Q*_c_(γ) for γ ≤ 0. The main difference from the *Q* = 0 behavior, is that there will be a concentration of the large abundances at roughly 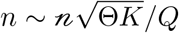 (instead of 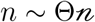 for *Q* = 0) since the peaks are suppressed by the niche interactions rather than the feedback from other types. This change in behavior will occur for 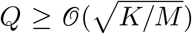. For the γ = 0 case as a function of *Q*, parallel work [81, 85] has explored the mainland model in detail. They find chaotic behavior consistent with our predictions and have developed numerical methods for analyzing the (simpler for γ = 0) dynamic mean-field equations. When the mainland model exhibits a diverse chaotic phase, our analysis strongly suggests that after some fraction of the types go globally extinct, the island model will exhibit the stable spatio-temporally chaotic phase. Roy et. al. [85] also show how spatial migration can stabilize chaos.

For positive γ the picture is murkier. In the mean-field framework of Equation 6, the feedback from responses of other types will be positive and this can destabilize the system by already abundant types becoming even more abundant. For *Q* = 0, on a single island the system will typically collapse to a small stable community. What happens with many islands we leave for future exploration.

Most analytical work for *Q* < *Q*_*c*_(γ) has focused on γ = 1 — perfectly symmetric interactions. In this special case, the Lagrange multiplier ϒ is a Lyapunov function which always increases in time and with stochastic population fluctuations the system is equivalent to conventional thermal statistical mechanics and “replica” methods can be used. With small migration from a mainland, recent work [32] has shown that there are exponentially many distinct stable states each with 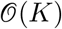 types surviving. However the system never really reaches any of these: it keeps wandering between higher and higher order saddles of the “landscape” of ϒ({*n*_*i*_}) more and more slowly, but with types going from populations of order *N/K* to order 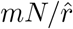. With an island model with perfectly symmetric *V*, it is not clear whether migration will lead to a desynchronized steadily slowing phase, or the system be unstable to local extinctions leading to global extinctions: but this should be analyzable by the present methods. A natural conjecture is that the slowing chaotic phase for γ = 1 — at least in the mainland model — is the limit of a stably chaotic phase as γ ↗ 1 with the equilibration time, *τ*_eq_ → ∞. This would be consistent if the chaotic phase exists for the whole 0 < *Q* < *Q*_*c*_(γ) plane for −1 < γ < 1.

## D.3 ASM mean-field analysis with fitness differences

Here we analyze the antisymmetric model with i.i.d. Gaussian distributed fitness differences {*s_i_*}, with variance 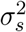. In particular, we wish to understand: how many types persist at the uninvadable fixed point? Note that the correct scale of comparison for the {*s_i_*} is the typical growth rate induced by the interactions, 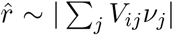. We know that 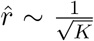 when Var[*V*_*ij*_] = 1; therefore the relevant parameter is the ratio 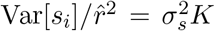. For 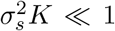 we expect that the fitness differences don’t much matter and half the types persist; for intermediate and large 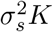, we can use mean field theory to predict the persisting fraction.

For convenience, we will carry out the analysis using the mean-field rescaling from Appendix B, so that the interactions contribute a total of *O*(1) to the growth rate, and the relevant parameter is simply 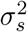. After carrying out the analyses, we will discuss what happens in the standard scaling.

In the mean-field rescaling, the mean-field dynamical equation is

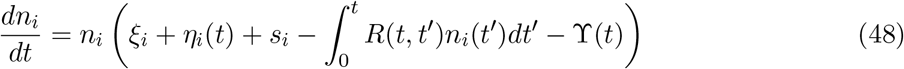

The Lagrange multiplier now attains a non-zero value due to the correlation between *s*_*i*_ and *n*_*i*_.

Time averaging the log rate of change gives us

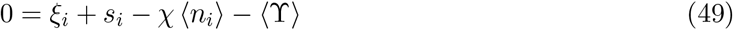

where again *χ* is the susceptibility 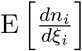. Note that the {*ξ*_*i*_} are Gaussian distributed with variance 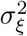. Solving for ⟨*n*_i_⟩, we have

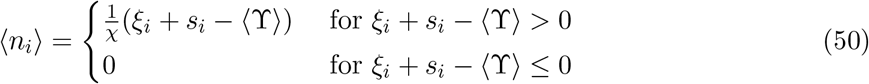

In order to determine what fraction of types persist, we must self-consistently solve for *χ*, 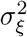, and 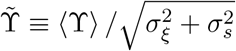 (rescaled for later convenience). Using the static moments of *n*_*i*_ averaged over the types/ensemble we have

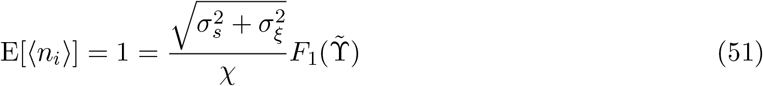

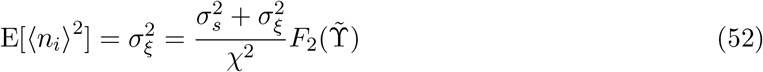

and from the definition of *χ*

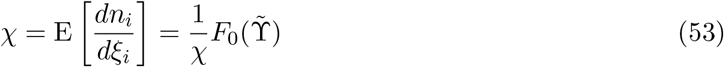

where we define the incomplete moment *F*_*m*_(*y*) as

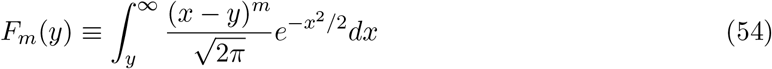

Note that 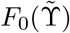 is the fraction of persistent types.

The system of equations can be most easily solved by eliminating *χ* and 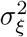 to obtain a non-linear equation for 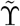, which can be easily solved for numerically. Starting with Equation 53, we have:

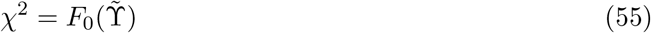

which then allows us to use Equations 51 and 52 to show

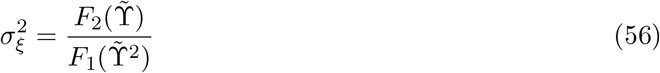

Finally, we arrive at a non-linear equation for 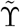 in terms of 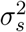:

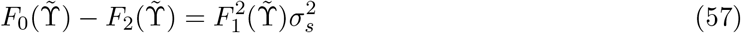

which can be solved for numerically.

In the limit of large 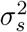, we can obtain some analytical understanding as well. Assuming dominant balance of the first two terms in Equation 57 we have:

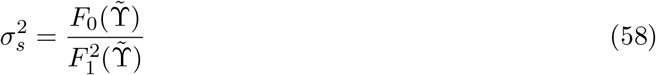

Making the ansatz of large ϒ, from the asymptotics of Gaussian integrals we have

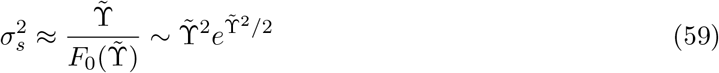

The leading order behavior is 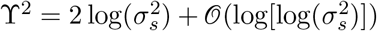, which gives us a persistent fraction 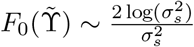.

The results of the analyses above describe the natural scaling if, as discussed above, the 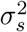 are replaced with 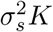. The fraction of persistent types goes as 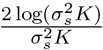 - meaning the total number of persistent types, 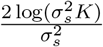, is increasing even for large 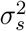. This can be confirmed numerically by computing the number of persistent at the fixed point for fixed 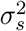, varying *K* (Figure 10, left panel). The mean-field prediction appears to hold quantitatively as well; plotting the persistent fraction as a function of 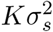 for various *K* and 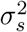 we get a universal curve, which is well matched by the numerical solution to Equation 57 (Figure 10, right panel).

**Figure 10:**
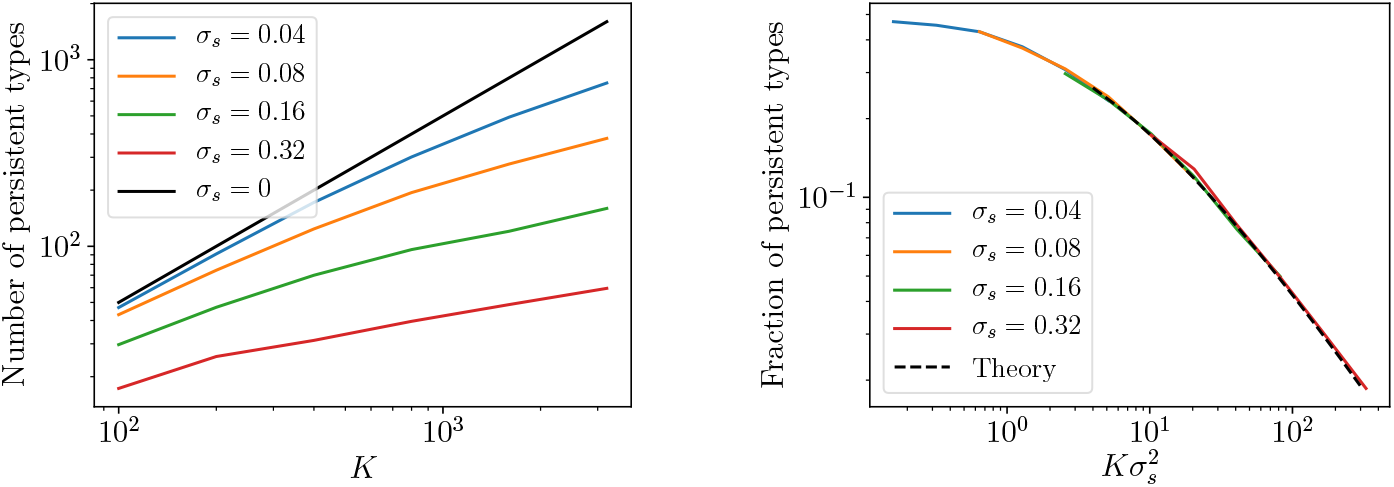
*Selective differences suppressing number of persistent types* for ASM. Fitness differences, *s*_*i*_, Gaussian distributed with variance 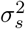. Number of persistent types increases, but logarithmically, with initial number, *K*, even for large 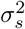 (left panel). Persistent fraction depends on 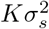 (left panel) and matches predicted scaling form from mean-field computation (right panel).

